# Organismal landscape of clock cells and circadian gene expression in *Drosophila*

**DOI:** 10.1101/2023.05.23.542009

**Authors:** Ines L. Patop, Ane Martin Anduaga, Ivana L. Bussi, M. Fernanda Ceriani, Sebastian Kadener

**Author notes:** To whom correspondence should be addressed Sebastian Kadener.

## Abstract

**Background:** Circadian rhythms time physiological and behavioral processes to 24-hour cycles. It is generally assumed that most cells contain self-sustained circadian clocks that drive circadian rhythms in gene expression that ultimately generating circadian rhythms in physiology. While those clocks supposedly act cell autonomously, current work suggests that in *Drosophila* some of them can be adjusted by the brain circadian pacemaker through neuropeptides, like the Pigment Dispersing Factor (PDF). Despite these findings and the ample knowledge of the molecular clockwork, it is still unknown how circadian gene expression in *Drosophila* is achieved across the body.

**Results:** Here, we used single-cell and bulk RNAseq data to identify cells within the fly that express core-clock components. Surprisingly, we found that less than a third of the cell types in the fly express core-clock genes. Moreover, we identified Lamina wild field (Lawf) and Ponx-neuro positive (Poxn) neurons as putative new circadian neurons. In addition, we found several cell types that do not express core clock components but are highly enriched for cyclically expressed mRNAs. Strikingly, these cell types express the PDF receptor (*Pdfr*), suggesting that PDF drives rhythmic gene expression in many cell types in flies. Other cell types express both core circadian clock components and *Pdfr*, suggesting that in these cells, PDF regulates the phase of rhythmic gene expression.

**Conclusions:** Together, our data suggest three different mechanisms generate cyclic daily gene expression in cells and tissues: canonical endogenous canonical molecular clock, PDF signaling-driven expression, or a combination of both.

## BACKGROUND

Most organisms use circadian clocks to keep temporal order and anticipate daily environmental changes. These clocks are based on self-sustaining biochemical oscillators that manifest at the molecular, physiological, and behavioral levels (1,2). Circadian clocks keep time through a complex transcriptional-translational feedback loop in the so-called “clock cells” (3,4). In *Drosophila*, the master transcriptional regulators CLOCK (CLK) and CYCLE (CYC) promote the transcription of several genes, thus activating the circadian system. Among them are *period* (*per*) and *timeless* (*tim*). The protein products of these genes are the main repressors of CLK-CYC mediated transcription, generating 24h mRNA oscillations (5). Post-transcriptional and post-translational modifications also play essential roles in circadian timekeeping (5–8). For example, both PER and TIM are post-translationally modified. The post-translational modifications of these proteins as well as the rates with which these modifications take place, significantly influence their activity and degradation rates and are key regulators of the circadian clock (5,9).

The most studied output of the circadian clock are the circadian rhythms in adult locomotor activity. In flies, this behavior is driven by approximately 75 circadian neurons *per* brain hemisphere. Circadian neurons are named after their physical location in the protocerebrum: Lateral Neurons (ventral, both small and large, and dorsal, sLN_v_, lLN_v_, and LN_d_, respectively), Dorsal Neurons (DN1-3), and Lateral Posterior Neurons (LPN) (3,10,11). While each clock neuron supposedly oscillates autonomously, the circadian neurons are organized into a neuronal network that is key for a coherent and robust behavioral output (1,12–16).

The Pigment Dispersing Factor (PDF) neuropeptide is the main neuromodulator of the circadian neuronal network in *Drosophila,* and it is produced by the small and large LNv (17). PDF expression is critical for the morning peak in locomotor activity in light:dark (LD) conditions and to sustain rhythmicity in constant darkness, DD (13,18,19). The PDF receptor is expressed in several brain regions as well as in some peripheral tissues and it is supposed to transmit the timing information to regulate the circadian behavioral outputs (12,16,20–26). However, PDF is not the only output neuropeptide of the circadian system. Other neuropeptides expressed in clock neurons such as neuropeptide F (NPF), short neuropeptide F (sNPF) and ion transport peptide (ITP) play a role in circadian system in *Drosophila* (28,29). Moreover, non-clock neurons transmit signals from pacemaker neurons to the circuits responsible for regulating patterns of movement and sleep-wake cycles through different neuropeptides such as diuretic hormone 44 (DH44) from the pars intercerebralis (PI) (30), or Leucokinin (LK) from the neurons that express the LK neuropeptide (31).

Circadian rhythms not only control behavioral rhythms but also a plethora of physiological processes across the body. These include many processes in peripheral tissues like the eye, the gut, the Malpighian tubules, and reproductive organs, all of which display daily rhythms in physiology (32–40). These rhythms include, among others, rhythms in eclosion (41), cuticle deposition (37), the release of cuticular pheromones by oenocytes (42), mating (43), and electrophysiological responses to odors in the antenna (38,44). Some of these rhythms are present in both LD and DD and hence they are self-sustained (the oscillations do not rely on signals from the environment), while others are robust in LD but dampen in DD. The general assumption is that rhythms in peripheral tissues are cell autonomous. Indeed, molecular assessments and reporter genes showed that core-clock components display oscillations in cells in several peripheral tissues (39,40,45–47). Moreover, microarray and RNAseq experiments showed that hundreds of mRNAs display daily oscillations in tissues like the eye, gut, Malpighian tubules, and fat body (48–50). However, it is unknown which cells within those tissues are responsible for the observed oscillations.

Peripheral circadian clocks can run independently from the clock in the brain or be modulated by it. Malpighian tubules (MT) are the paradigm of a tissue that is independent of circadian control by the central brain: PER oscillations in MT continue in decapitated flies (36), and the phase of TIM oscillation is maintained in DD when the MT is transplanted into the abdomen of flies entrained to antiphase LD cycles (34). Other peripheral clocks have their own molecular clock, but their oscillation phase can be modulated by PDF. This is the case of oenocytes or the prothoracic gland in the fly pupae which drives eclosion rhythms (46). In addition, the clock in the fat body exhibits *per* oscillations in LD but requires a functional clock in the brain for displaying DD oscillations (51). The visual system encompasses both peripheral tissues (eye) and the central nervous system (optic lobe) and it is responsible for transmitting at least part of the photic information to the brain (11). In the retina, visual pigment, sensitivity, and rhabdomere size cycle daily in LD and DD (52,53). In the Lamina, L1 and L2 interneurons and the glial cells that enwrap each lamina cartridge change their size in a circadian fashion (54–58). However, L2 have constant levels of *per* and high levels of *Pdfr,* suggesting their rhythms are driven by PDF (59). While these results suggest different modes of regulation of target tissues by the circadian clock, this has not been systematically tested.

Here, we aim to tackle this question: how is circadian gene expression achieved and regulated in different tissues across *Drosophila*? To do so, we first identified which *Drosophila* cells express core-clock components by employing single-cell and bulk RNAseq datasets. We then utilized circadian datasets from different tissues in combination with single-cell sequencing data to identify cell types expressing mRNAs that are known to cycle. Using these approaches, we identified new sets of neurons expressing core-clock genes in the *Drosophila* brain. In addition, we found that a third of the cell types in the fly body express core-clock genes. Interestingly, many cells lacking clock gene expression display high PDF receptor (*Pdfr*) levels. Some of these cell types express high levels of mRNAs that present a circadian pattern, strongly suggesting that PDF drives daily gene expression in these cell types. Other cell types in peripheral tissues express circadian clock components and *Pdfr*, suggesting that PDF might influence their rhythmic gene expression but it is not required to generate these rhythms. Together, our findings suggest that while most cell types exhibit daily mRNA cycling, they achieve so through three distinct mechanisms.

## RESULTS

### Using standardized expression of core-clock components to identify cells harboring a circadian clock

We used publicly available single-cell data from the brain, the optic lobe, and the whole fly to identify cell types that harbor a circadian clock in the fly body (60–63). First, we mapped the expression of the main core-clock gene components: *tim, per, Clk, cyc, cry, cwo,* and *vri,* focusing primarily on the expression of *tim*, *per*, and *Clk* given that *vri* and *cwo* have clock-independent functions (64–67) and *cyc* mRNA is known to be widely expressed (68).

Due to the characteristic low input of single-cell RNA sequencing techniques, it can be challenging to determine with certainty that a given cell is not expressing a specific mRNA, especially for low-expression genes such as *Clk*. As core-clock components display significant changes in abundance during the day, this dropout problem could be even more profound: we might not detect a gene because of the sample acquisition time instead of because that gene it is not expressed in a particular cell type. As there is no information about the dissection time of each tissue, this is an intrinsic limitation of mapping mRNAs that display daily changes in abundance in these datasets. To overcome these limitations, we focused on analyzing the abundance of genes by cell type rather than single cells. We achieved this by adding the signal from all the cells of the same type, which should mitigate this dropout problem. Additionally, we identified clock cells based on expression of genes that cycle with opposite phases (*cry* and *Clk* vs. *tim*, and *per*) to diminish the possibility of missing core-clock expressing cells due to the dissection time (Figure 1A, Supplementary Table 1). Moreover, *tim* mRNA levels are high, which allows identification of core-clock expressing cell types even if not dissected at the optimal timepoint.

**Figure 1:**
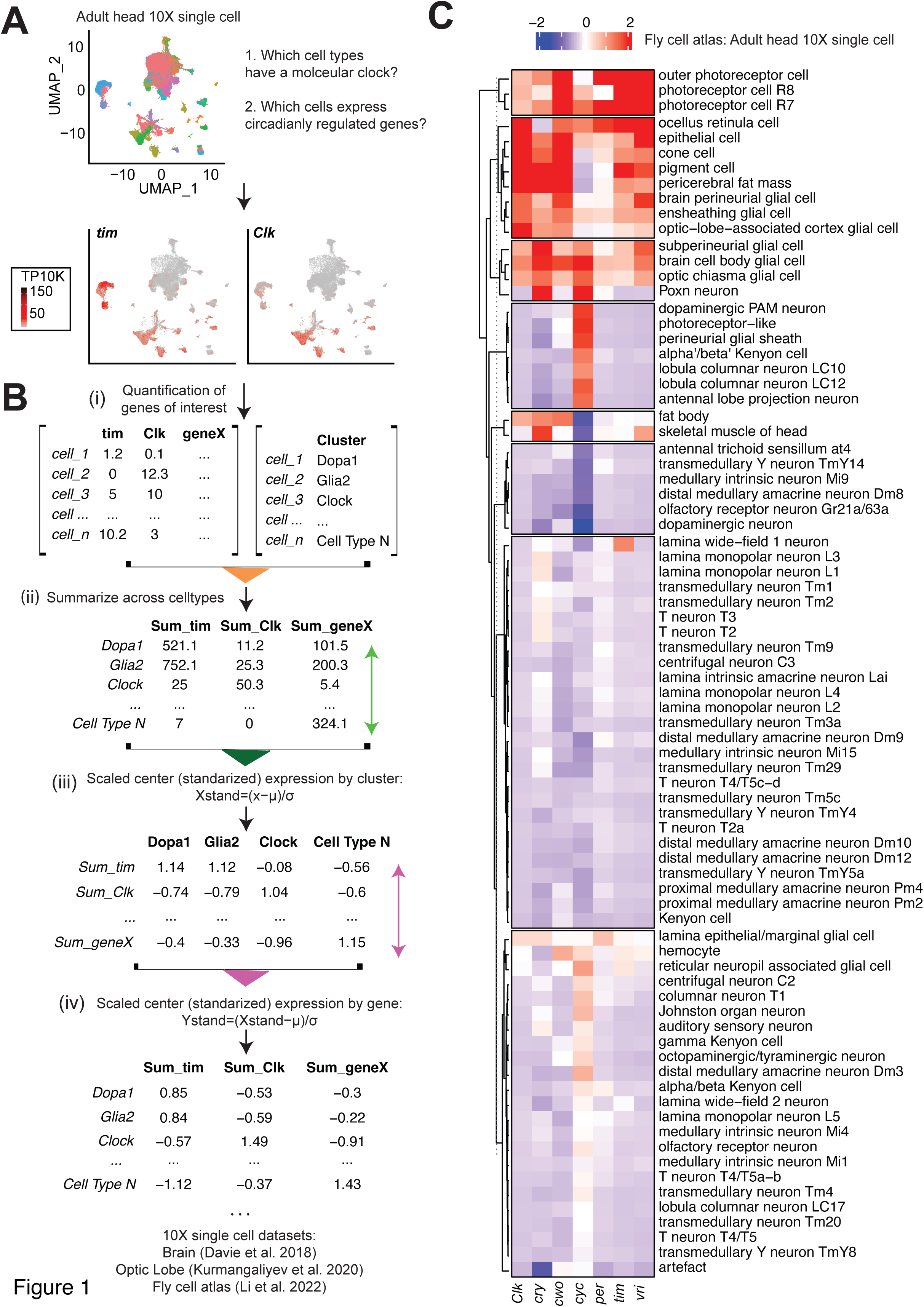
Standardized gene expression recapitulates known core-clock expression pattern in the head. **A.** Representative UMAP plot from the fly head single cell dataset colored by cell type assignment, and showing *tim* and *Clk* gene expression. Red indicates higher expression (color bar, TP10K). **B.** Representation of the processing pipeline used for analysis: i) Generate the matrix of raw counts of gene expression *per* cell type; ii) Calculate the sum expression of each gene by cell type; iii) Standardize the expression values across clusters; iv) Standardize those expression values across genes. We performed this analysis for the following 10X single cell datasets: Brain (Davie et al. 2018), Optic Lobe (Kurmangaliyev et al. 2020), and Fly cell atlas (Li et al. 2022). **C**. Heatmap of standardized values of core-clock genes from Head 10X data with blue and red representing low and high levels, respectively. Clusters are calculated with k means (data: Li et al. 2022).

To identify cells with core-clock gene expression, we standardized the gene expression value across cell types and genes (Figure 1A-B, Supplementary Table 1). Briefly, after extracting the normalized gene expression levels (Figure 1B i) and adding up the signal for each mRNA *per* cluster (Figure 1B ii), we calculated how much each gene contributes to the transcriptome of each cell type. We achieved this by normalizing and centering gene expression across each cell type (Figure 1B iii). Finally, we compared the relative contribution of each clock gene between different cell types by normalizing and centering all gene expressions across clock genes (Figure 1B iv). This approach has the advantage of considering the variation between genes and between clusters. To validate this approach, we used it to identify which cell types in the fly head express clock genes (Figure 1B, Supplementary table 1). As expected, we found that core-clock genes are highly enriched in photoreceptors and glial clusters. We explored the raw normalized reads and verified that we could recapitulate the general single-cell signal in each cluster (Supplementary Figure 1A). We also verified these results by analyzing a bulk RNA-sequencing dataset from sorted fly head cells (69) (Supplementary Figure 1B). Importantly, this dataset includes sorted LN_v_s, in which we found enrichment for *Pdf* and clock genes. Therefore, we concluded that the standardized expression allows the identification of cells harboring a circadian clock even in single-cell data.

### Only a fraction of cells in the fly body expresses detectable levels of core-clock genes

We then extended our analysis to the rest of the fly body, starting with data from the entire body wall (Figure 2A-B). This dataset contains many cell types, but only oenocytes, epidermal and epithelial cells, fat cells, and hemocytes are enriched for at least two core-clock genes. Indeed, some of these cell types (epithelial, tracheal, glial, and fat cells) generally express clock genes in most of the inspected tissues (Figure 2C-H and Supplementary Figure 2 and Supplementary table 1). Interestingly, the expression of core-clock genes in the muscle varies among tissues. While the muscle in the antenna expresses most clock genes (Supplementary Figure 3A), other muscles in the body seem to be enriched only for *vri* (body wall), *cry* (heart), or *vri* and *cry* (leg, wing, and haltere) and likely do not have a circadian clock (Figure 2B-H). To ensure that we do not miss circadian gene expression in these muscle cells (*i.e.*, these cells might support core-clock gene expression at lower levels than other cells within the same tissue), we checked the co-expression of *tim* (the highest expressed clock gene) and *Mhc* (a muscle cell marker) in muscles of different tissues (Supplementary Figure 2A). We utilized *tim,* as it is highly expressed and generally is possible to detect it even at the low timepoints. Indeed, we found that most cells with high *Mhc* levels have none or very low expression of *tim*. We validated these results by analyzing total RNA sequencing from mature flight muscles (70). This dataset indicated that clock genes such as *tim, Clk, vri,* and *per* might be present in some muscle cells but at low levels (Supplementary figure 2B). In sum, our results suggest that some muscle cells express clock genes and others do not, suggesting that core-clock gene expression in the muscle cells happens only in some organs in flies.

**Figure 2:**
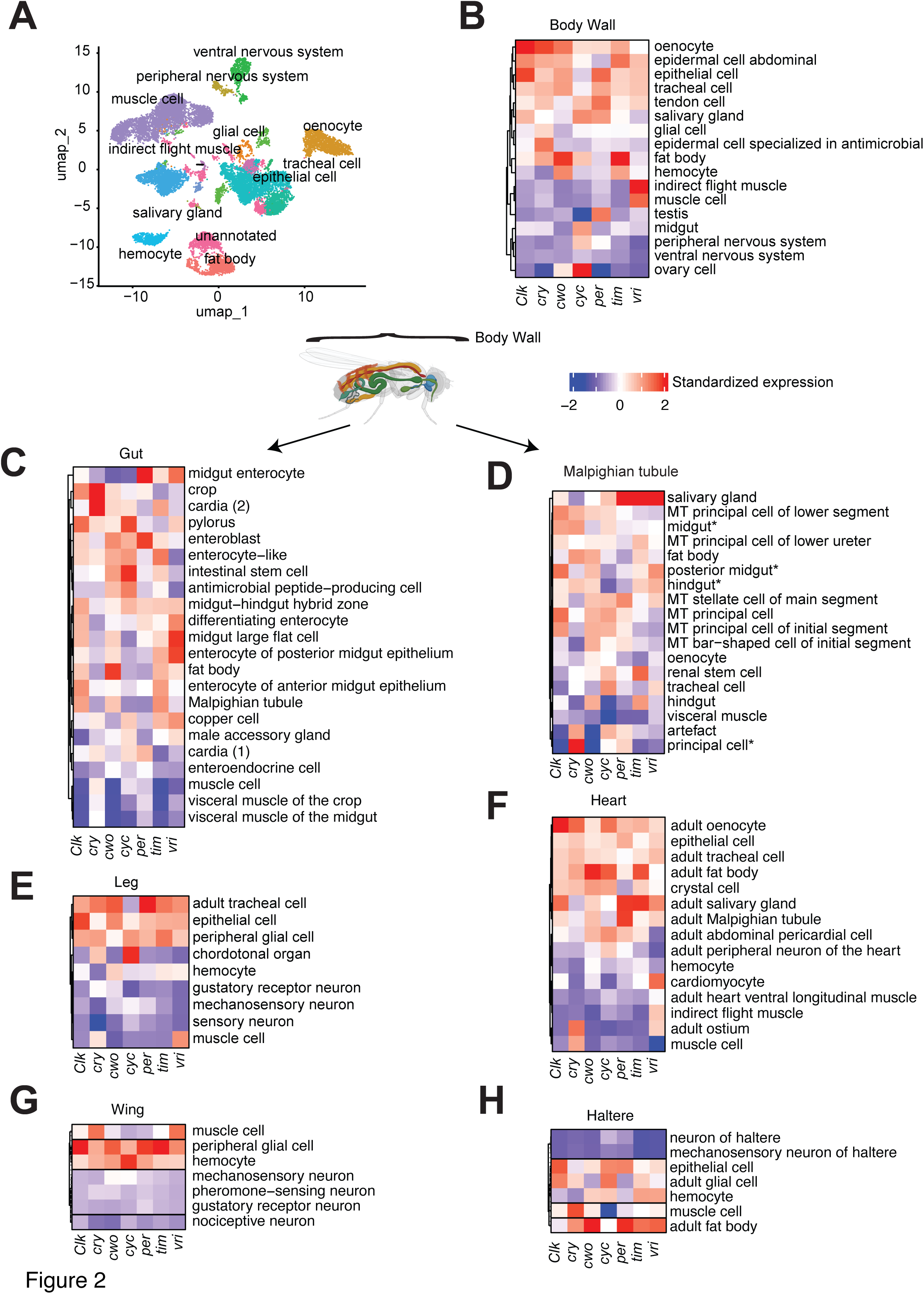
Cell types across the body have diverse core-clock enrichment patterns. **A**. Representative UMAP of body wall 10X single-cell data colored by cell type assignement. **B-H**. Heatmaps of standardized values of core-clock genes from 10X data of the body wall, gut, Malpighian tubule, leg, heart, wing, and haltere in *Drosophila*. Blue and red represent low and high expression levels, respectively. Clusters are calculated with k means (data: Li et al. 2022).

Most cell types in the Malpighian tubules and a large fraction in the gut display enrichment for core-clock genes, supporting the idea that they have a functional clock (Figure 2C-D). The exceptions are the visceral muscle cells and the enteroendocrine cells. On the other hand, only some cell types in the wing, leg, heart, and haltere express core-clock genes (Figure 2E-H). Surprisingly, core-clock genes are not enriched in neurons in these body parts. As in the rest of the body, in these different organs, the expression of clock genes is observed mainly in epithelial, tracheal, and glial cells.

Surprisingly, we found no expression of clock genes in the olfactory receptor neurons (ORN) in the antenna or in gustatory receptor neurons (GRN) in the proboscis (Supplementary Figure 3A-B). Given prior reports that show cycling physiology in the ORNs and GRNs (33,38,44,71), we further examined clock-gene expression levels in ORNs and GRNs by looking at their expression at the single-cell level (Supplementary Figure 3C-D). To exclude the possibility of a data processing artifact, we used the available visualization tool from the Aerts lab (SCope, (60)). Briefly, we inspected the co-expression of *tim, elav* (neuronal marker), and *Orco* (olfactory receptor neuron marker) in these data. We found almost no overlap between neurons (marked by *elav* expression) and *tim* (Supplementary Figure 3E-F), except for the photoreceptor neurons in which we also found expression of clock genes (Supplementary Figure 3E). These data indicate that most neurons in the brain and body do not harbor a functional circadian clock and that photoreceptors and clock neurons constitute the exception rather than the rule.

Last, we analyzed core-clock enrichment in reproductive tissues. We first focused on the testis, in which previous work had shown that the lower testes epithelium and the seminal vesicles express *Clk, tim,* and *per* in a circadian manner (32). In the male reproductive gland, we found high enrichment for core-clock genes in the seminal vesicle, ejaculatory bulb, and duct (Supplementary Figure 4A). Within this organ, we also found enrichment in somatic cells like the epithelial cells, pigment cells of the reproductive tract and the cyst (Supplementary Figure 4B). On the other hand, in ovaries, *tim* and *per* are highly expressed in the follicle cells but excluded from oocytes and nurse cells (72). Our results recapitulated these observations (Supplementary Figure 4C). However, the roles of TIM and PER in this tissue are not circadian-related (72). Interestingly, both ovaries and testis have cells with highly enriched *cyc* levels (i.e., some spermatocytes and germ cells).

### Enrichment analysis in the brain identifies known and novel clock-gene-expressing cell types

Surprised by the fact that most neurons in peripheral tissues express low level of core-clock genes, we continued by characterizing the enrichment levels of these genes using single-cell data from the fly brain (60). First, we characterized the total contribution of each cell type to the bulk signal for each core-clock gene in the whole brain (Supplementary Table 2). As expected, glial cells are the main contributor of core-clock gene expression. Among the glial cells, the ensheathing glia (which accounts for 6% of the brain cells in the single-cell dataset) is the major contributor (23% of the total *tim* signal). Interestingly, cell clusters that are much smaller, like clock neurons and Lawf 1 and 2 cells (see below), are also major contributors to *Clk* and *tim* expression, respectively. Circadian neurons contribute 10% of the total *Clk* signal in the dataset despite representing only about 0.4% of the cells (Supplementary table 2).

We then performed the analysis using the standardized expression for each gene by cluster, as described in the previous section (Figure 3 and Supplementary Figure 5). As expected, we found *Pdf* neuropeptide expression restricted to the central pacemaker clock neurons (a cluster that includes the LN_v_ neurons), and we found core-clock genes were strongly enriched in clock neurons and glial cells (73,74) (Figure 3A and Supplementary Figure 5). Interestingly, we found lower levels of *cwo* among some glial cell types (astrocytes, ensheathing glia, and a single unannotated glial cluster), suggesting that in these glial subtypes, the circadian clock does not utilize *cwo* (Supplementary Figure 5). Not surprisingly, as previously shown in the head dataset (Figure 1), we also found high expression of clock genes in cell types known to express circadian clock components, like photoreceptors (75), or that display circadian physiological changes like hemocytes (76) (Figure 3). Additionally, some poorly-defined cell types, *i.e.,* a group of cholinergic clusters in the central brain and optic lobe (Vesicular acetylcholine transporter positive neurons VAChT +), have enrichment for *cwo* and *cyc* (Supplementary Figure 5A). This is not surprising given the wider non-circadian related expression of those genes.

**Figure 3:**
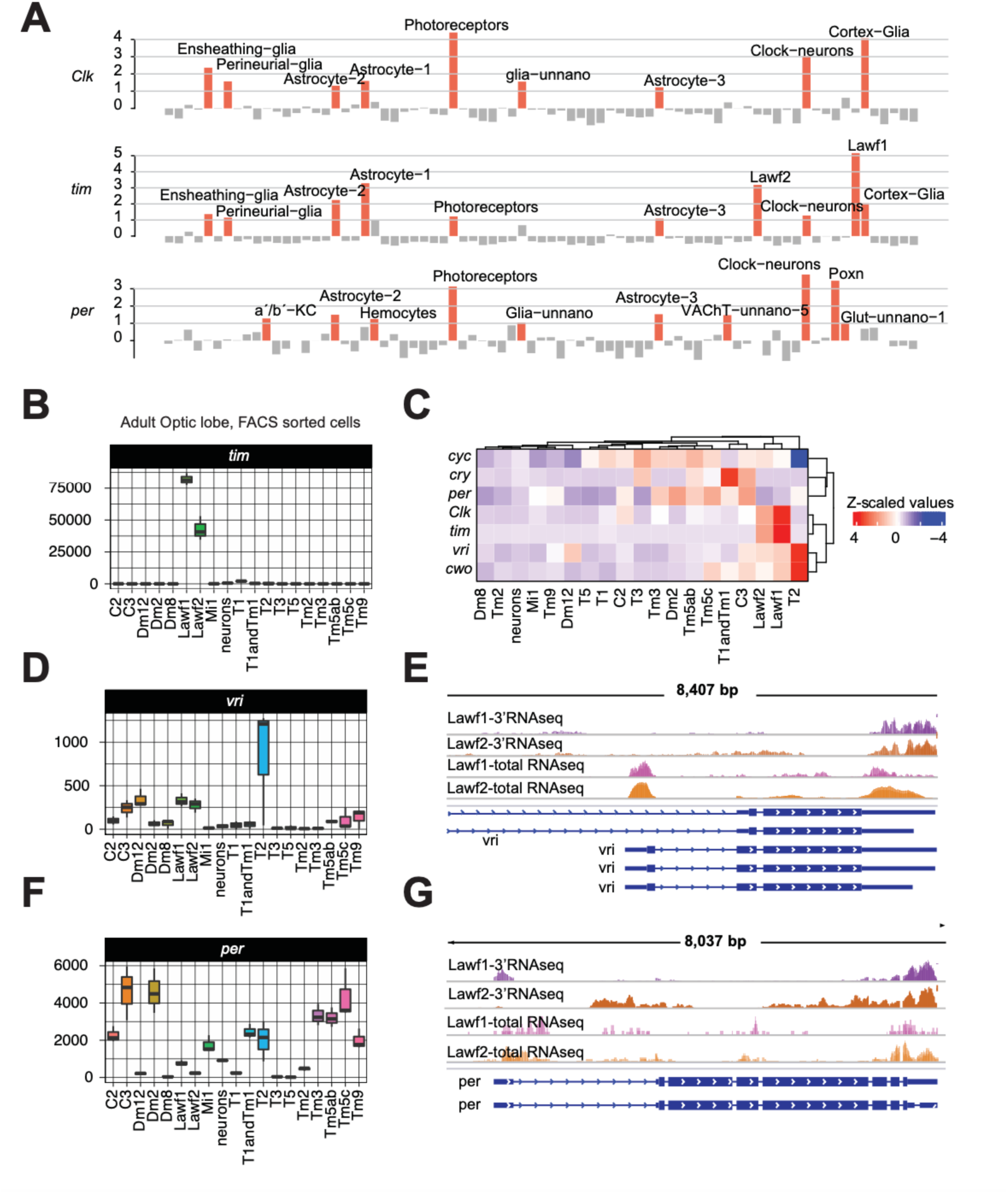
Core-clock gene enrichment analysis in the brain reveals novel cell types expressing high levels of core-clock components. **A**. Standardized core-clock gene expression from 10X single-cell data from the brain. In red, cells with expression levels higher than one standard deviation from the mean (data: Davie et al. 2018). **B.** Boxplot showing normalized *tim* expression in FACS sorted cells from the optic lobe (data: Konstantinides et al. 2018). **C** Heatmap of scaled normalized expression of core-clock genes in FACS-sorted cells from the optic lobe (data: Konstantinides et al. 2018). **D.** Boxplot showing normalized *vri* expression in FACS-sorted cells from the optic lobe. **E.** IGV snapshot of *vri* expression in total RNA and 3’RNA sequencing from sorted Lawf1 and Lawf2 neurons (data: Konstantinides et al. 2018 and Davis et al. 2020) **F.** Boxplot showing normalized *per* expression in FACS-sorted cells from the optic lobe (data: Konstantinides et al. 2018). **G.** IGV snapshot of *per* expression in total RNA and 3’RNA sequencing from sorted Lawf1 and Lawf2 neurons (data: Konstantinides et al. 2018).

Interestingly, we detected high levels of some core clock genes in three new neuronal groups: Pox-neuro positive neurons (Poxn) and Lamina wide field neurons type 1 and 2 (Lawf1 and Lawf2 neurons) (Supplementary Figure 5A). Poxn cells are particularly enriched for *per* (at levels comparable to the clock neurons), and both Lawf1 and Lawf2 clusters have the highest enrichment for *tim* in the fly brain (even higher than the clock neurons; Figure 3A). We corroborated the enrichment of these genes in the Lawf1 and Lawf2 cells using a different single-cell dataset (62) (Supplementary Figure 5B) as well as additional optic lobe RNA sequencing data from single cells and sorted cell populations (61,69) (Figure 3B-G). These datasets confirmed a high expression of *tim* in both Lawf1 and Lawf2 neurons (Figure 3B). Additionally, the sequencing depth of these data allowed us to assess the presence of less abundant core-clock genes (Figure 3C), confirming the expression of *vri* and *per* in these cell types (Figure 3D-G). Notably, the Lawf cells only display signal from the most proximal *vri* promoter, which is utilized by the isoform required for circadian rhythms (64). In addition, we also found low but detectable *per* and *vri* expression in the C2, C3, and T2 neurons using the FACS sorted dataset (Figure 3D-G). Regarding *Clk* expression, even though it seems particularly enriched in Lawf1 and Lawf2 cells (Figure 3C), the levels are too low to be 100% certain that this gene is expressed in all of them (approximately 100 reads, Supplementary table 3). If there is indeed cycling in these cell types, these low *Clk* levels could be due to the dissection time (when *tim* expression is high, *Clk* expression is low). Alternatively, its expression could be restricted to a subset of these cells.

The lamina wide-field feedback neurons subtype 2 (Lawf2) are relevant for orientation to visual cues and, thus far, have not been ascribed to any circadian function. Their cell bodies are located in the medulla cortex and project towards the medulla and lamina neuropils (77–79). To explore the possibility that Lawf2 cells support a functional circadian clock, we employed a characterized Lawf2 driver (80) to drive the expression of a green fluorescent reporter (GFP) to localize them within the medulla cortex of the optic lobe along with a *tim* fluorescent transcriptional reporter previously characterized (12). PDF staining was included to add a location reference for the Lawf2 cells. We found that Lawf2 somas and their lateral projections are located in the vicinity of PDF^+^ large ventral Lateral Neurons (l-LN_v_s) projections (Figure 4A). As predicted by the transcriptomic data, a proportion of the GFP^+^ Lawf2 cells also express *tim* as also observed in the single cell RNAseq data (Figure 4B). Given that the oscillation of *tim* abundance is a hallmark of a functional molecular clock, we assessed the immunoreactivity of the *tim* reporter in Lawf2 cells across a day in flies maintained under 12:12 light-dark conditions. To do so we utilized our *tim-*TOMATO transcriptional reporter, that as we previously showed has a peak of expression in the late night/early morning (12). Indeed, we found that the levels of the *tim* transcriptional reporter change as a function of the time of day both in general and, in particular, in the Lawf2 cells suggesting that the Lawf2 cells might indeed harbor a functional circadian clock (Figure 4C-D).

**Figure 4:**
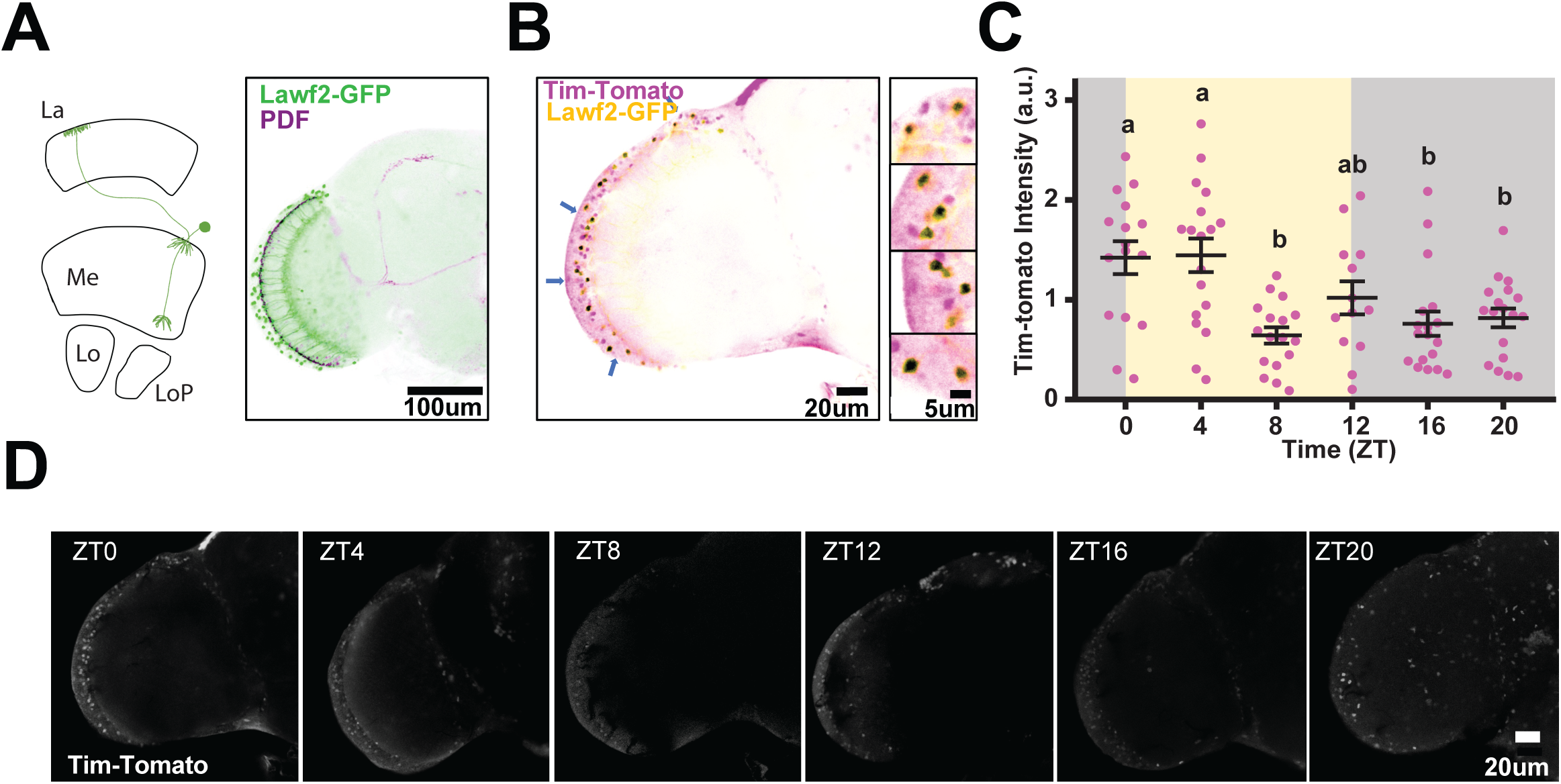
*tim*-Tomato transcriptional reporter shows high levels of *tim* in Lawf2 cells. **A.** Represenation of Lamina wide field neurons and their projections in the optic lobe (four stratified neuropils: the lamina (La), medulla (Me), lobula (Lo), and lobula plate (LP) and Lawf2-driven GFP expression in *w^1118^*; UAS-GFP; Lawf2-Gal4 fly brain immunostained with anti-GFP (green) and anti-PDF (purple) antibody. The regions where both signals colocalize are marked in black. **B.** *w^1118^*; UAS-GFP/*tim*-Tomato; Lawf2-Gal4 fly brain immunostained with anti-GFP (yellow) and anti-dsRed (magenta) antibodies. Blue arrows indicate the cells that are shown in magnified photos on the right. The regions where both signals overlap are marked in black. **C**. *tim*-Tomato intensity quantification in Lawf2-GFP expressing cells across time points. ZT (zeitgeber time) being ZT0 and ZT12 the time at when lights turn ON and OFF, respectively. Each dot on the plot represents the mean intensity value for a single brain. ZT0 = 1.422 + 0.165 (n=16), ZT4 = 1.445 + 0.169 (n=18), ZT8 = 0.643 (n=17) + 0.082, ZT12 = 1.019 + 0.165 (n=13), ZT16 = 0.760 + 0.122 (n=18), ZT20 = 0.818 + 0.094 (n=18); N=2. One way ANOVA followed by Bonferroni’s multiple comparisons test, F(5,94) = 6.691, p< 0.0001. Only different letters (but not the combination of them) indicate significant differences: p<0.05 for ZT0 vs ZT16, ZT0 vs ZT20, and ZT4 vs ZT20; p<0.01 for ZT4 vs ZT16, and ZT0 vs ZT8; p<0.001 for ZT4 vs ZT8. Data expressed in Mean+SEM. **D.** Representative images of *tim*-Tomato brains immunostained with anti-dsRed antibody dissected at the indicated time points.

### mRNAs displaying daily changes in expression are enriched in cells without the molecular circadian clockwork

Previous works have identified hundreds of genes expressed in a circadian manner in fly heads and brains (48,81–86). We generated a new polyA bulk RNA sequencing dataset from brains of *w^1118^* flies dissected every four hours, which allowed us to identify almost 300 strong cyclers in the *Drosophila* brain (Supplementary Table 4), of which more approximadetely 200 hundred display strong signal in the single cell sequencing data. To understand where these genes are cycling within the brain and whether the circadian clock controls these genes directly or indirectly, we utilized an available single-cell dataset to identify which cell types express these cycling genes (60).

We found that cycling genes are highly expressed in many cell types within the fly brain (Figure 5A). Specifically, we determined that around 50% of the cycling genes are enriched in the glial cell, hemocytes, and photoreceptors (Figure 5A). This is not unexpected as the glia accounts for more than 90 % of the gene expression of core-clock genes within the brain (Supplementary table 2). The remaining 50% are present in a combination of cells in the brain. Given that many of these genes are being expressed in cell types without a known endogenous circadian clock, we explored different options for their potential expression. We are aware that the presence of the expression of these genes that present circadian oscillation does not mean that they are being expressed rhythmically in that cell type. However, many of these cycling genes are highly enriched in the same cell populations. In this context, their rhythmic expression must result from these cell populations 1) having an endogenous clock or 2) being controlled by the central pacemaker cells through specific(s) neurotransmitter(s).

**Figure 5:**
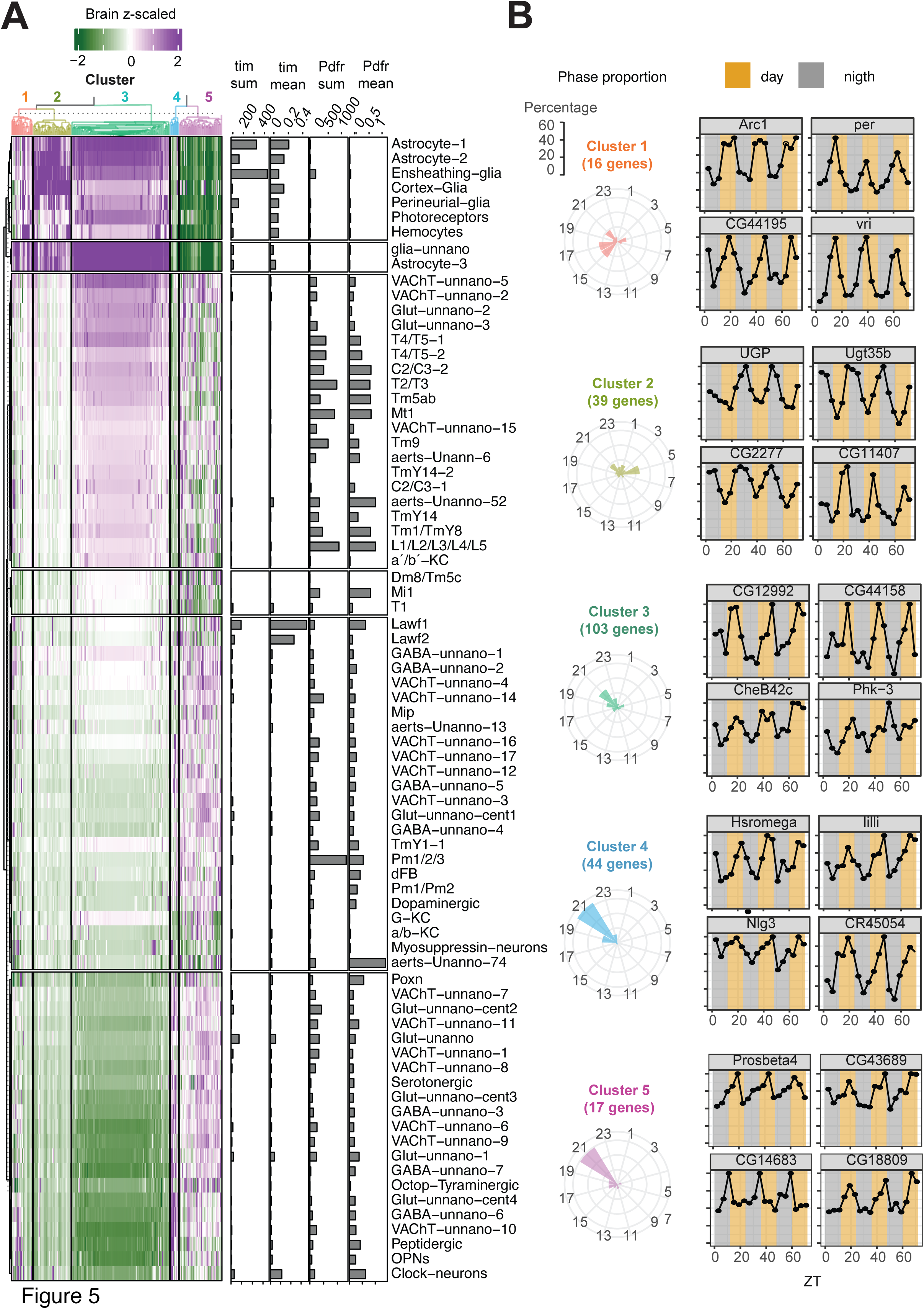
Genes with cyclic expression in the brain are enriched in cells with or without core-clock expression. **A**. Enrichment of cyclic genes in single-cell fly brain clusters. Heatmap of standardized expression values clustered by similarity. In bar plots, sum and mean expression of *tim* and *Pdfr* for each cluster. **B**. Normalize expression of representative genes of each cluster across 3 days (12-hour/12-hour LD). (right) and proportion of genes by cluster cycling at each phase (left). Expression is normalized by the maximum (n=3, 30 flies per replica per timepoint, FDR<0.05 and amplitude > 1.5) and phase is calculated using JTK algorithm.

As PDF is the most studied neurotransmitter of the clock, we decided to evaluate the expression of *tim* and the PDF receptor (*Pdfr*) as proxy for defining groups of cells that harbor a self-sustained clock and those that can potentially be regulated through PDF respectively. We utilized both the Sum and Mean expression values of these genes in the different neuronal clusters in order to get a fair representation of their expression levels and prevent biased conclusions. For example, if we focus only on the Sum, we would miss the *tim* expression in the Lawf2 cells due to the low number of cells in this cluster. On the other hand, if we were to focus only on the Mean, we would missed one of the clusters in the ensheathing glia that expresses *tim* (Figure 5A). Unsurprisingly, we found enrichment for *tim* in photoreceptors and several glial cell types. Strikingly, we found that most of the other cell types expressing genes that cycle (and don’t express core clock components) are enriched for *Pdfr* (Figure 5A, lower three panels).

To further understand the different brain cycling genes, we performed cluster analysis of the mRNAs that display cyclic expression. We found that the majority of the oscillating mRNAs cluster into five different clusters and investigated the expression of genes in each of the groups (Figure 5B). We observed that the phase distribution of the genes varies across each cluster. Interestingly, cycling genes enriched in cells with high levels of *Pdfr* (clusters 4 and 5) have mostly a peak at 21 hours (Figure 5B) suggesting their oscillations are regulated by PDF.

We then expanded this analysis to the peripheral tissues. As many cycling genes are tissue-specific, we focused on the gut, and Malpighian tubules, the only tissues for which temporal RNA sequencing data across the day is available (48,50). Litovchenko *et al*. (2021) found hundreds of genes displaying 24 h rhythms in gene expression in each of those tissues (Figure 6A). Similar to what we observed in the brain, the expression of genes with circadian oscillation in these tissues is highly cell-type specific (Figure 6B-C). Furthermore, some identified cyclers are heavily enriched in cells without a circadian clock, such as the visceral muscle cells in the gut (Figure 6B). Importantly, visceral muscles present high levels of *Pdfr* (Figure 6B). Finally, most cycling genes in the Malpighian tubules (MT) originate from MT principal cells, which have core-clock gene expression (Figure 6C). In sum, we found that the expression of genes that oscillate in each of the tested tissues correlates with the expression of *tim* or *Pdfr*.

**Figure 6:**
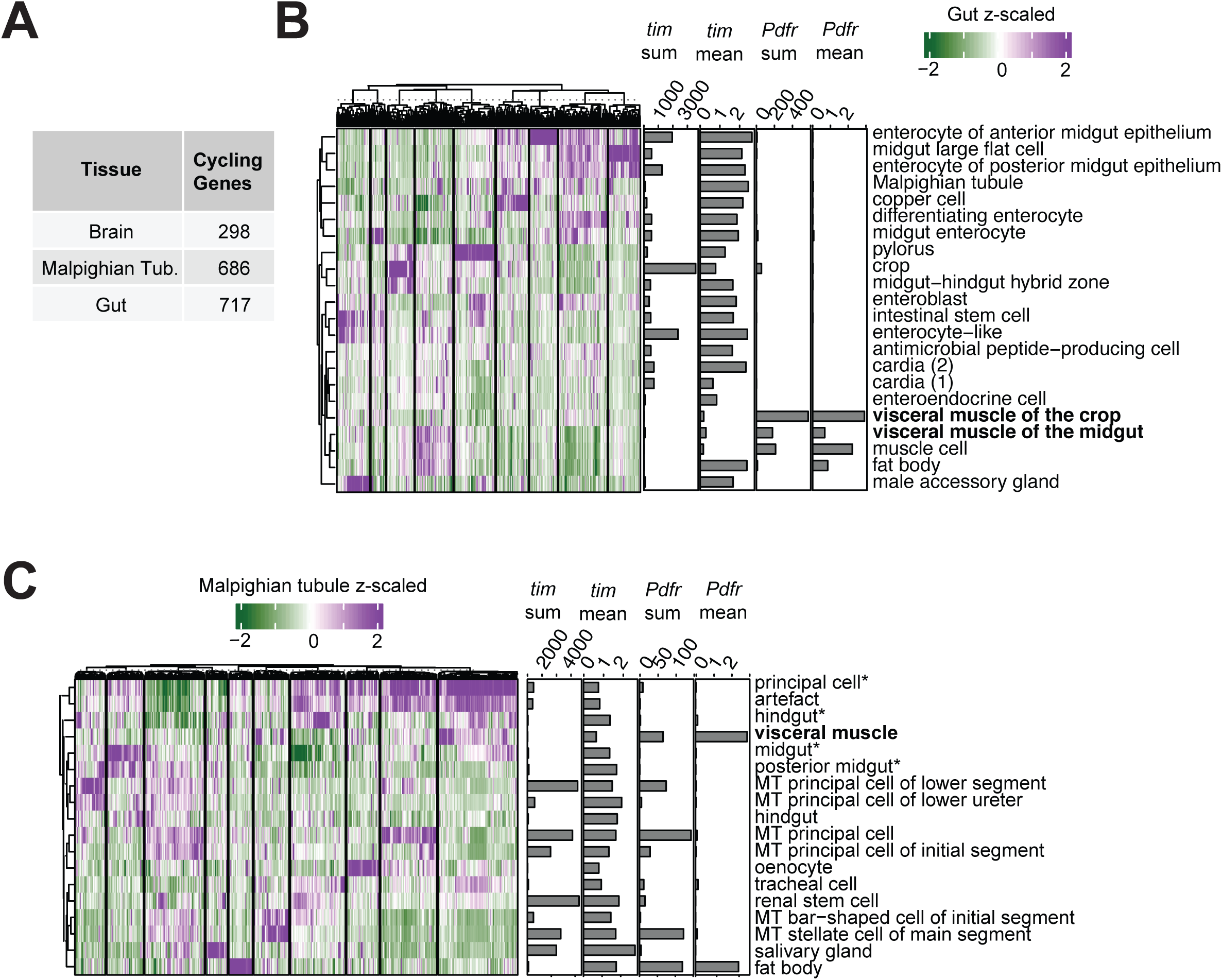
Expression of cycling genes in peripheral tissues correlate with high levels of core clock genes or *Pdfr*. **A**. Table summarizing the number of cycling genes identified in each tissue from Litovchenko *et al.* 2021. **B.** Heatmap displaying enrichment of genes that cycle in the gut in single-cell clusters. **C.** Heatmap of enrichment of cyclic genes in the Malpighian tubule in single-cell clusters. In bar plot, sum and mean expression of *tim* and *Pdfr* for each cluster.

### Circadian gene expression profiles might be achieved through three different pathways

The presence of genes with cyclic gene expression in cells without a known circadian oscillator but with high expression of the PDF receptor suggests that mapping the presence of *Pdfr* could provide insight into the potential regulation of peripheral tissues and cell types by the central pacemaker. To this end, and given that certain cell types enriched for core-clock genes are present in multiple tissues (i.e., hemocytes, epithelial, and glial cells), we decided to integrate the scaled analysis from all the different single-cell datasets analyzed in this work based on the expression of the core clock gens and the *Pdfr* (60,62,63). Integrating all the scaled data avoids potential issues due to differences in their sequencing deepness. We then clustered all the cells from the different datasets analyzed in this manuscript (Figure 7A). Importantly, some cell types were present in more than one dataset and cluster together, demonstrating the accuracy of this integration. For example, the Lawf and Poxn neurons from the brain and head datasets cluster together. Something similar happens with the epithelial cells: they cluster within the same group independently of the tissue of origin.

**Figure 7.**
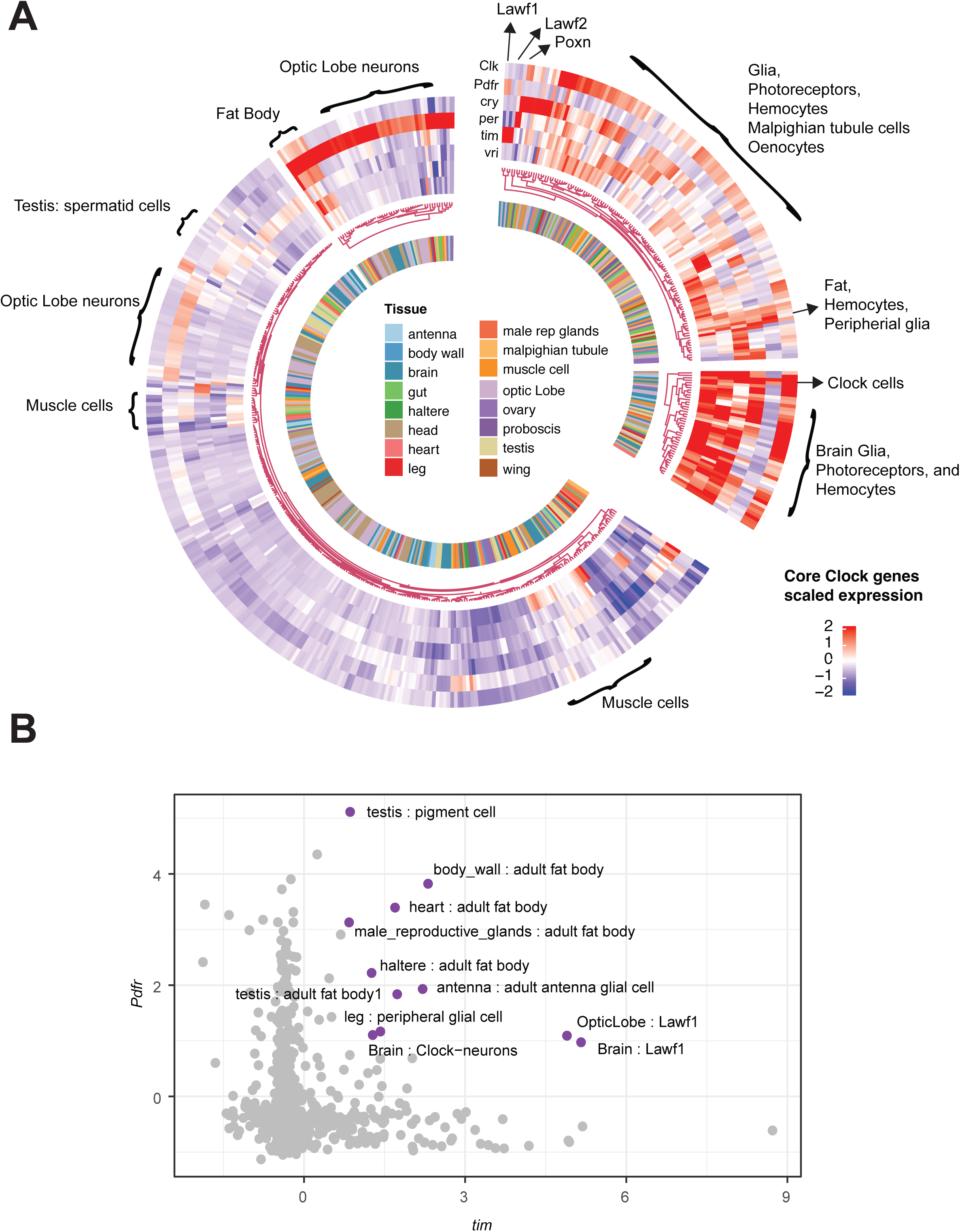
Expression of *Clk*, *per*, *tim*, *vri*, and *cyc* core clock genes and *Pdfr* across the whole *Drosophila* body suggest different patterns of circadian regulation. **A**. Heatmap of standardized expression values clustered by similarity. Blue and red represent decreased or increased expression, respectively. The colors in the inner part represent the tissue of origin for each cluster. Main cell types of interest are identified individually. **B.** Standardized expression of *tim vs. Pdfr.* In violet, the clusters with high *tim* and *Pdfr* scaled expression.

Having validated the integration of the different single-cell datasets, we characterized the overlap between core clock genes and *Pdfr* expression in each cell type. We immediately appreciated the presence of cells with very high and low enrichment levels for these genes (Figure 7A, right vs. left sections). In particular, we can classify the cell types into four different groups according to their relative enrichment of core clock genes and *Pdfr*.

First, we found a fraction of cells that are enriched for both core clock genes and *Pdfr*. Under this category, we found circadian neurons, fat cells originating from different tissues, hemocytes, epithelia, and peripheral glial cells. Indeed, fat body cells are the cells with the highest enrichment for *Pdfr* in the whole body (Figure 7A, Supplementary table 1). Second, we identified cells enriched for core clock genes but not for *Pdfr*. This group is comprised mainly of glial cells from the central brain, epithelial cells, all Malpighian tubule cells, and oenocytes (Figure 7A, Supplementary table 1). Among these, we also recognize cells that are particularly enriched for *tim* (Lawf neurons), *per* (Poxn neurons), or *vri* (muscle cells) (Figure 4.1 A, Supplementary table 1). Third, we detected cells highly enriched for *Pdfr* but not for core clock genes. Most of these cells are part of the optic lobe, such as L1/2/3, Dm, or Tm neurons (Figure 7A, Supplementary table 1). Finally, we found that nearly 50% of the cells were not particularly enriched for either *Pdfr* or any clock gene (Figure 7A, Supplementary table 1). This percentage can be an overstatement as we are measuring enrichment compared to the rest of the cells of the same dissected tissue.

With the exception of the fourth group described above, the cell types express core clock components or *Pdfr.* We then focused on comparing expression of *tim* (the highest expressed core clock component) and *Pdfr* in all the cell types described in this study. Strikingly, we found that cells express *tim* or *Pdfr* but not both genes together (Figure 7B, Supplementary table 1). There are few cells that express both *tim* and *Pdft* at high levels including certain glial cells, fat cells, and clock neurons (all of which are in the first group described above). Interestingly, Lawf1 cells are also part of this exception, although they express *tim* at much higher levels than *Pdfr*.

Together, these results highlight three different mechanisms in which daily gene oscillations can be generated on the fly: 1) cells can have a molecular clock independent of the central pacemaker neurons (*i.e.* Malpighian tubule cells), 2) cells can have a molecular clock regulated by the central pacemaker neurons by PDF (*i.e.* fat body cells) and 3) cells can have oscillations driven by PDF (*i.e.* optic lobe neurons and some muscle cells across the body) (Figure 8).

**Figure 8:**
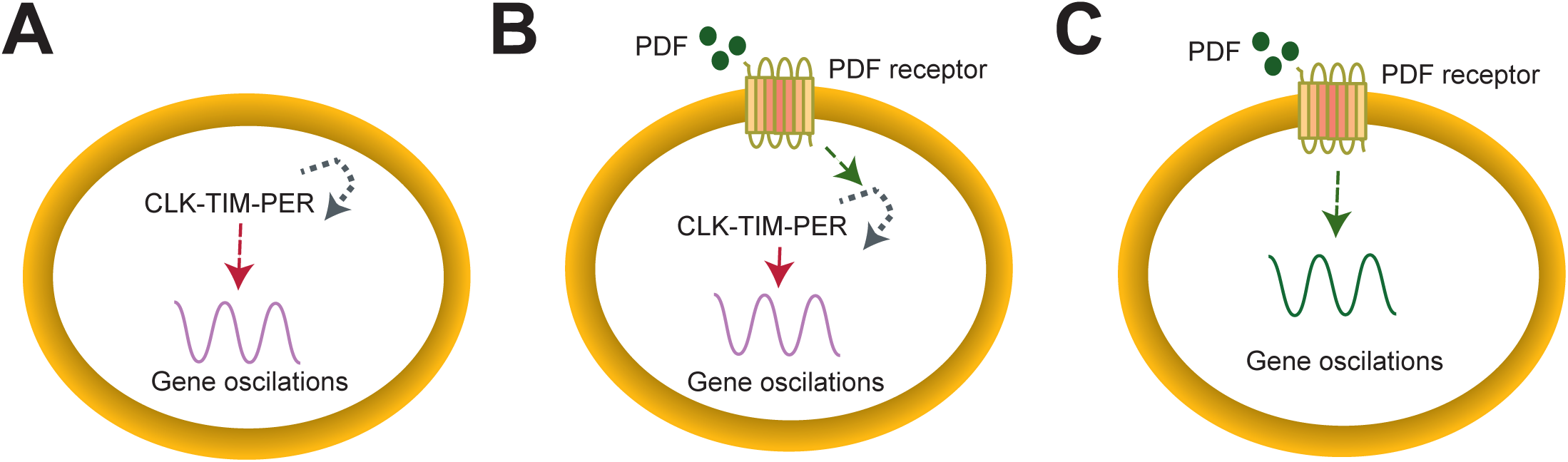
Three different possible models of circadian gene regulation. **A**. Cells have an endogenous clock that is independent of PDF **B.** Cells have an endogenous molecular clock that can be tuned by PDF **C.** Cells do not have an internal molecular clock but express high levels of *Pdfr*, suggesting that the central pacemaker could regulate observed oscillations in gene expression through PDF signaling.

## DISCUSSION

In this work, we mapped the expression of core clock genes in the body of *Drosophila melanogaster* using a standardized expression of core clock genes. This method helps get a general picture of each tissue but has clear limitations in situations where a gene might be expressed at low levels. Notably, the datasets analyzed do not include information about the circadian time of collection of the samples. Consequently, this analysis could have missed cell types with circadian expression because of sampling time. We tried to overcome this limitation by examining groups of genes such as *tim-per-vri vs Clk-cry*, which cycle with opposite phases enabling us to detect core-clock gene expression regardless of the collection time of the sample. Despite these limitations, this approach allowed us to recapitulate expected core-clock expression in all tissues with verified circadian pacemakers like clock neurons, fat body, glial cells, photoreceptors, and Malpighian tubules. Interestingly, we found enrichment for clock genes in tracheal, glial, fat, epithelial cells, and hemocytes across many tissues. While we are the first to describe the existence of a circadian clock in these tissues across the fly body, analog cells in mammals (lung epithelia, macrophages, and other epithelial cells) harbor a circadian clock (87–91).

However, some observations challenge previous reports in other cell types. For example, we could not detect core clock genes enriched in neuronal clusters in the antenna, proboscis, or maxillary palps. This lack of core clock machinery in the Olfactory Receptor Neurons (ORN) and Gustatory Receptor Neurons (GRN) is surprising, as they display circadian changes in physiology that are abolished by the expression of a dominant negative allele of *Clk* (33,44,71). The observed discrepancy could be due to several factors: 1) only a small fraction of the GRN and ORN express core-clock genes, and they were not detectable in the single-cell datasets 2) these cells do not harbor a circadian clock and are regulated remotely by circadian neurons. Experiments with cell-type resolution monitoring RNA and protein levels at different timepoints will be key to solving this potential discrepancy. Similarly, early work using a *per-*GFP reporter demonstrated circadian oscillations in the leg and wing and showed florescent signal in the basal cell of sensory bristles and chemosensory cells of the wing margin (40). However, our analysis shows enrichment primarily in epithelial and glial cells in these tissues.

Interestingly, our work identified new potential clock neurons: the Poxn and Lawf1/2 neurons. Specifically, we found that Poxn neurons express high levels of *per* while Lawf1 and 2 express the highest levels of *tim* mRNA in the fly brain. In the case of Lawf neurons, we validated these results using additional bulk and single-cell RNA sequencing datasets and a *tim* transcriptional reporter. Bulk RNAseq data show low but detectable levels of other clock genes like *vri* and *per* in Lawf neurons. This could indicate that these cells could potentially have a functional circadian clock. Indeed, given the generally low expression of *Clk* mRNA, it would seem more likely that most, if not all, Lawf cells express all circadian components. So far, we corroborated the cyclic expression of *tim* (highest in the early morning) in Lawf2 by ICC using our previously established *tim*-Tomato reporter line.

Finally, we mapped the expression of known 24-hour cyclers across their available single-cell datasets. We found that only around 50% of these cyclers were enriched in cells with a reported circadian clock, with the remaining genes enriched in cells with no clock gene expression. Of course, the fact that they are expressed in those cell types does not mean that their expression is cyclic. However, many of these cycling genes are so highly enriched in the cell populations, and their added signal accounts for a significant fraction of the total signal in the brain making unlikely that the observed cycling in whole brains comes from anywhere else. The finding that some cells without a circadian oscillator could express a subset of mRNAs that oscillate daily suggests that temporal information is conveyed through communication with cells with a functional circadian clock. This could be achieved through electrical or neurohormonal communication. As PDF is the main neuropeptide produced and used by the central circadian pacemaker cells (12,16,20–25,92), we mapped the presence of receptors for this neuropeptide in the fly head and peripheral tissues.

Based on the relative enrichment of core clock genes and *Pdfr*, we classified the cell types on the *Drosophila* body into four groups (Figure 7, 8). The first group is comprised of cells that are enriched for core-clock genes and *Pdfr*. This is a heterogeneous group. One the one hand, it includes the clock neurons, which are self-sustained but use PDF for synchronizing their circadian gene expression increasing the robustness of the system. In addition, this groups includes other cells groups that likely use PDF to adjust the phase of the oscillations in response to signals from the circadian neurons (Figure 8B). The latter includes multiple fat cells (from the body wall, heart, haltere and antennae), some hemocytes, and peripheral glial cells (legs and antennae). The second group includes cells enriched only for core-clock genes but that do not express *Pdfr* (Figure 8A). Within this group are glial cells, Malpighian tubule cells, and oenocytes. These are cells that either contain a self-sustain circadian oscillator that cycles robustly in DD independently of the central pacemaker (Malpighian tubes, (36)) or cells that maintain cycling after several days under free-running conditions (day 6 in DD) like oenocytes. The phase of cycling in the oenocytes has been shown to be altered in *Pdf* mutants, but as these cells lack *Pdfr,* this is likely an indirect effect from the pacemaker cells (oenocytes, (46)). In this second group of cells, we also found cell types particularly enriched for *tim* (Lawf neurons), *per* (Poxn neurons), or *vri* (muscle cells) (Figure 7A). The fact that one component of the clock is predominant (while the others are at low levels) might indicate a non-circadian function for those genes in these cells or the presence of a different non-canonical regulation. Third, we detected a group of cells in which the *Pdfr* is highly expressed, but there is no core-clock gene expression (Figure 8C). In the brain, most of these cells are part of the optic Lobe cells, such as L1/2/3, Dm, or Tm neurons (Figure 7A). Previous work using a *Pdfr* reporter found expression in the Lamina, demonstrating that the observed expression is not an artifact of the data processing (93). Moreover, previous work in L2 showed that these cells expressed high levels of *Pdfr* but low, not-cycling, *per* levels (59). Cells belonging to this group could receive information from the central pacemaker through the neuropeptide PDF, which could lead to the generation of oscillations of subsets of mRNAs. Alternatively, some of these cell types could be directly regulated by the Lawf1/2 cells that synapse with several of these groups. Other additional cells in the body also belong to this category, including the visceral muscle in several tissues (Figure 7). Interestingly, previous work showed that overexpressing *Pdfr* in the visceral muscles in a *Pdfr* mutant fly recover PDF-induced renal motility (94). Based on our results, it will be interesting to determine if the circadian oscillations in visceral muscle of the gut are fully dependent on the *Pdfr*, as those cells do not express core clock components. Finally, we also found that about 50% of the cell types do not show enrichment for core-clock genes *nor Pdfr*, particularly neurons from the antennae, optic lobe, and central brain.

Classical work in the field assumes that circadian gene expression is consequence of cell autonomous presence of the circadian clock on those cells. However, our data shows that different cell types utilize different strategies to achieve daily cyclic gene expression: (1) cells that have a molecular clock independent of the central pacemaker neurons (e.g. Malpighian tubule cells); (2) cells that have a molecular clock regulated by the central pacemaker neurons by PDF (e.g. oenocyte, fat cells across the body and the clock neurons themselves) and (3) cells in which their oscillations are driven by PDF (e.g. gut visceral muscles and optic lobe neurons) (Figure 8). Importantly, PDF is not the only known circadian neuromodulator (31,51,95,96). Indeed, recent work have shown the existence of several circuits that convey information from circadian neurons to other circuits within the brain using other neurotransmitters as Leucokinin (LK), Neuropeptide F (NPF), and short NPF (sNPF). Therefore, we might have underestimated the central regulation of peripheral cells by the central pacemaker cells, and the community will benefit from exploring the expression of additional neuropeptide receptors using additional datasets. In this context, it is likely that the circadian neurons regulate the timing in gene expression in a large fraction of the brain and body through expression of different neuropeptides.

### CONCLUSIONS

Our work established for the first time the enrichment of core-clock genes and 24-hour cyclers at the cell type level in the whole fly. This allowed us to identify new potential circadian cells as well as to get new insights into how timing information is transmitted from the brain pacemaker cells to other cells in the body. By looking for cells in the brain expressing core clock genes, we found that Lawf and Poxn cells are enriched for *tim* and *per*, respectively. Furthermore, we verified that some Lawf2 cells express *tim* and that their *tim* level fluctuates around the day by immunostaining.

In addition, by analyzing the cell-type enrichment of cyclically expressed mRNAs in brain and peripheral tissues, we identified oscillating genes enriched in cells that do not express core-clock genes but have high levels of *Pdfr.* Together, our results suggest that three different mechanisms generate daily gene expression oscillations: an internal molecular clock independent of PDF signaling, an internal clock tuned by PDF, or a PDF-driven expression (Figure 8).

## MATERIALS AND METHODS

### Fly husbandry

*w^1118^* and UAS-mcD8-GFP (referred to as UAS-GFP) fly strains were obtained from the Bloomington Drosophila Stock Center (Indiana, USA). Lawf2-Gal4 line (11D03-Gal4) were gifted by Dr. Claude Desplan (80). Lawf2-Gal4 lines were recombined with the UAS-GFP line to generate the *w^1118^*; UAS-GFP; Lawf2-Gal4 line. *tim*-Tomato reporter flies consist of a codon-optimized and destabilized tdTomato fluorescent protein, which has three copies of a nuclear localization signal (NLS) to focus signals for quantification under the control of the *tim* promoter (12). All crosses were performed and raised at 25 °C in a 12:12 Light/Dark cycle (LD) conditions.

### Immunohistochemistry

*w^1118^*; UAS-GFP; Lawf2-Gal4 line was crossed to the *tim*-Tomato reporter line. +; UAS-GFP/*tim*-Tomato; Lawf2-Gal4) males were selected from their progeny. Three to five days old flies were anesthetized with CO_2_, their whole body fixed in 4% PFA for 1 hour at room temperature (RT), and their brains were dissected in PBS at ZT0, ZT4, ZT8, ZT12, ZT16, and ZT20. Brains were subjected to a standard immunostaining protocol. Briefly, the brains were incubated ON at 4 °C in a mix of mouse anti-GFP (1:1000, Sigma) and rabbit anti-DsRed (1:1000, Rockland) and 7% normal goat serum in PBS. Samples were washed in PBS and incubated with anti-mouse Alexa 488 (1:500, Jackson ImmunoResearch) and anti-rabbit Cy3 (1:500, Jackson ImmunoResearch) for 2 hours at RT, washed in PBS and mounted in anti-fade mounting medium Fluoromount-G (Southern Biotech).

Single snapshots for each brain were obtained using a Zeiss LSM 880 confocal microscope. Image analysis was performed in Fiji (ImageJ, NIH-USA). Statistical analysis and data visualization was performed in Prism 8 (GraphPad).

#### Generation of RNAseq datasets

*w^1118^* flies were entrained for at least 3 days at 25 °C and dissected at six different timepoints (ZT3, ZT7, ZT11, ZT15, ZT19, and ZT23). RNA from the fly brains was extracted using TRIzol reagent (Sigma, T9424) and treated with DnaseI (NEB, M0303L). We used 150 ng of RNA as input for preparing 3’ RNA sequencing libraries following CelSeq2 protocol (97,98), changing the UMI to 6 bases. Sequencing was performed on Illumina NextSeq 500 system.

#### Circadian gene assessment from bulk RNA sequencing

For the 3′ RNA-seq, data were aligned to the *Drosophila melanogaster* dm6 genome and transcriptome version using STAR (99), and quantification was done with ESAT (100). The circadian analysis was performed using the package MetaCycle (101). For each circadian timepoint, three replicates were analyzed. Since each replicate was sequenced separately, the counts were divided by the maximum value in each replicate after normalizing by library size. Genes with more than two zero counts at any time point were discarded. The amplitude for each replicate was calculated as the maximum divided by the minimum for each gene. The JTK algorithm was used for the circadian analysis (102). A gene was considered a cycler if the JTK p-adjusted value was less than 0.05 and the amplitude was more than 1.5.

#### Single-cell RNA processing

Single-cell pre-processed data was downloaded from the respective GEO databases. For the fly cell atlas (63), an in-house script was written to integrate the data into Seurat format (https://github.com/ipatop/FlyCellAtlas.download.with.annotations). Briefly, we gathered the data and performed the normalization and clustering necessary to access the complete list of normalized genes. Cell type assignment was extracted from the published data. We validated the cell-type assignment by manually inspecting marker genes.

The fly cell atlas datasets come from dissected tissues except for the fat body, coenocyte, and tracheal cells that were FACS sorted using a tissue-specific GAL4 driving UAS–nuclear-GFP (Li et al 2021).

For the optic lobe dataset (62), we downloaded the data from GEO (GEO: GSE156455). Integration and clustering steps were needed to process this dataset. We used the parameters reported by the authors (62).

For enrichment and clustering analysis, gene expression was retrieved using the *FetchData* function from Seurat. Gene level sum was summarized by cell type using general R functions. After that, z-scaling and centering were done first over the gene level and then over the cell type level. Heatmaps and clustering were done using the ComplexHeatmap package. The optimal k-mean cluster number was determined using the *eclust* function with 500 randomization steps. All functions used for this analysis are stored in the following R-Package: https://github.com/ipatop/SingleCell_SumScaledExpresion.git

#### Sorted cell-type RNA processing

Data was downloaded from GEO (Series GSE116969 (69), and GEO: GSE103772 (61)) and aligned to the *Drosophila melanogaster* dm6 genome and transcriptome version using STAR. Quantification was done with feature counts. Mean normalized counts were used with ComplexHeatmap for clustering and visualization. The optimal k-mean cluster number was determined using *eclust* function with 500 randomization steps.

## DECLARATIONS

### Ethics approval and consent to participate

Not applicable

### Consent for publication

Not applicable

### Availability of data and materials

The datasets generated and/or analyzed during the current study are available in the GEO repository. The data generated in this study has been deposited in GEO (accessing number GSE233184).

## Funding

This work was funded by the NIH R01 grant (R01GM125859) to SK.

### Authors’ Contributions

ILP performed the computational analysis, contributed to the conceptualization of the work and writing the manuscript; AMA contributed to the conceptualization of the work, designed and performed the circadian experiments, sample collection, RNA sequencing, and edited the manuscript; ILV performed the immunohistochemistry experiments and analysis, FC designed the immunohistochemistry experiments and edited the manuscript; SK designed the experiments, conceptualized the work, and wrote the manuscript.

## Acknowledgments

We thank Dr. Claude Desplan for the Lawf2-Gal4 fly lines and Dr. Sagiv Shifman for his insight on the data analysis processing.

## Authors’ information

One or more of the authors of this paper self-identifies as an underrepresented ethnic minority in science. One or more of the authors of this paper self-identifies as a member of the LGBTQ+ community.

## SUPLEMENTARY FIGURES

**Figure S1.**
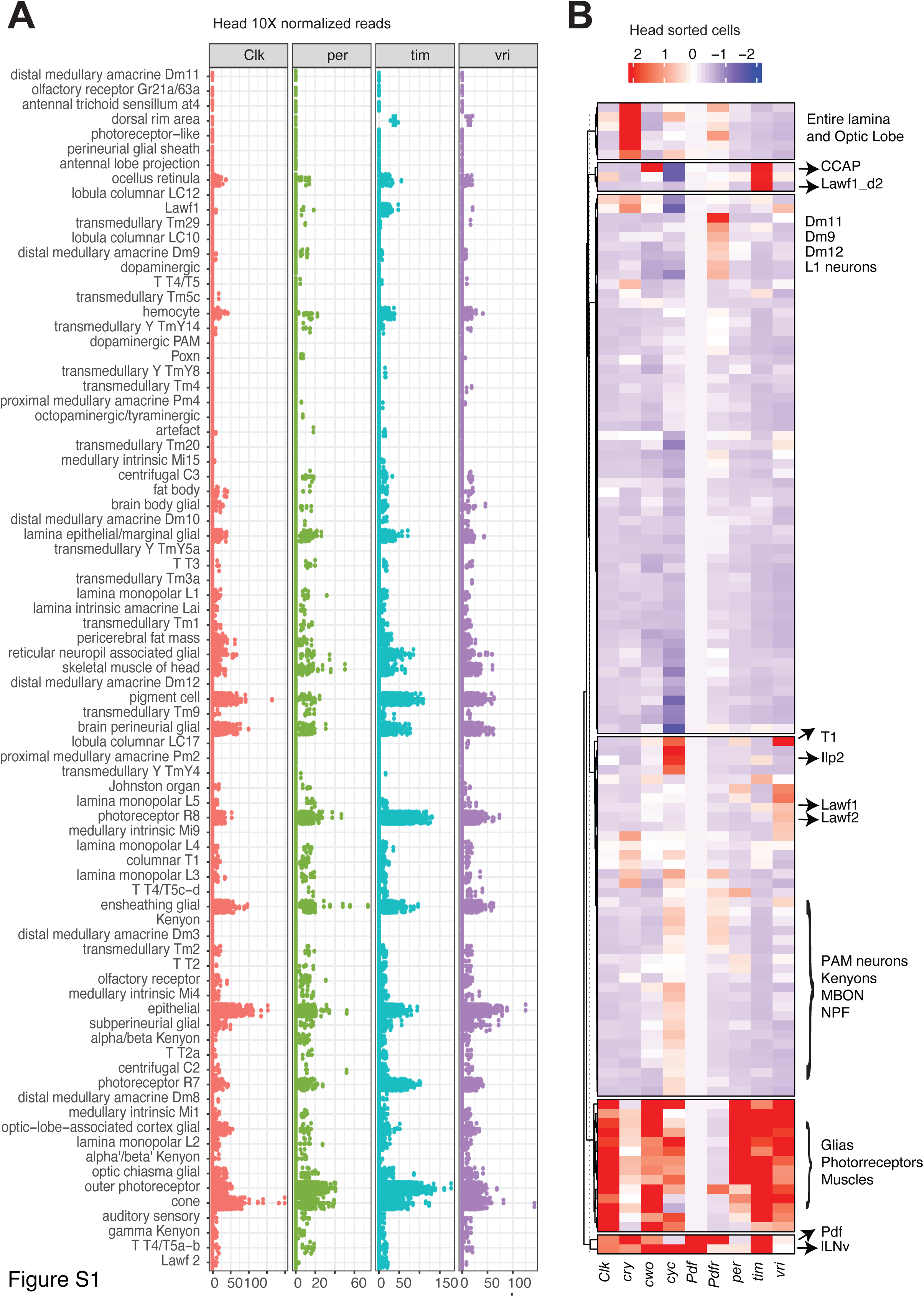
(related to figure 1): Standardized gene expression recapitulates known core-clock expression pattern in the head. **A.** Dotplot of normalized gene expression in each cell by cluster. Each dot is a cell. **B.** Heatmap of z-scaled mean values with blue and red representing low and high levels of sorted cells followed by bulk RNA sequencing.

**Figure S2.**
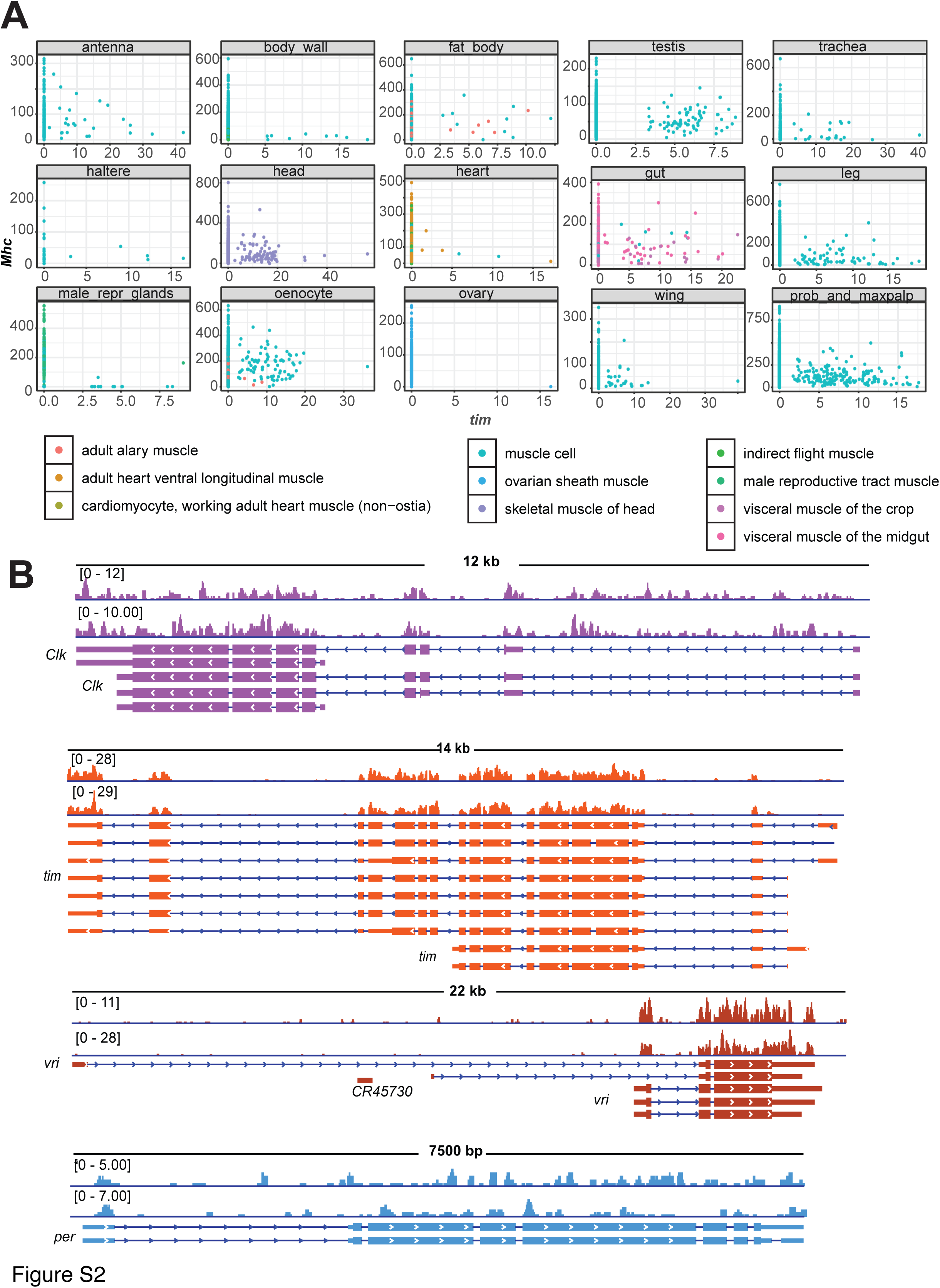
(related to figure 2): Muscle cells express low levels of core-clock genes. **A.** Dotplot of normalized gene expression *tim vs. Mhc*. Each dot is a cell. Colors represent cell type. Data is separated into boxes according to tissue origin. **B.** IGV snapshot of *Clk, tim, vri,* and *per* expression in total RNA sequencing from dissected mature flight muscle.

**Figure S3.**
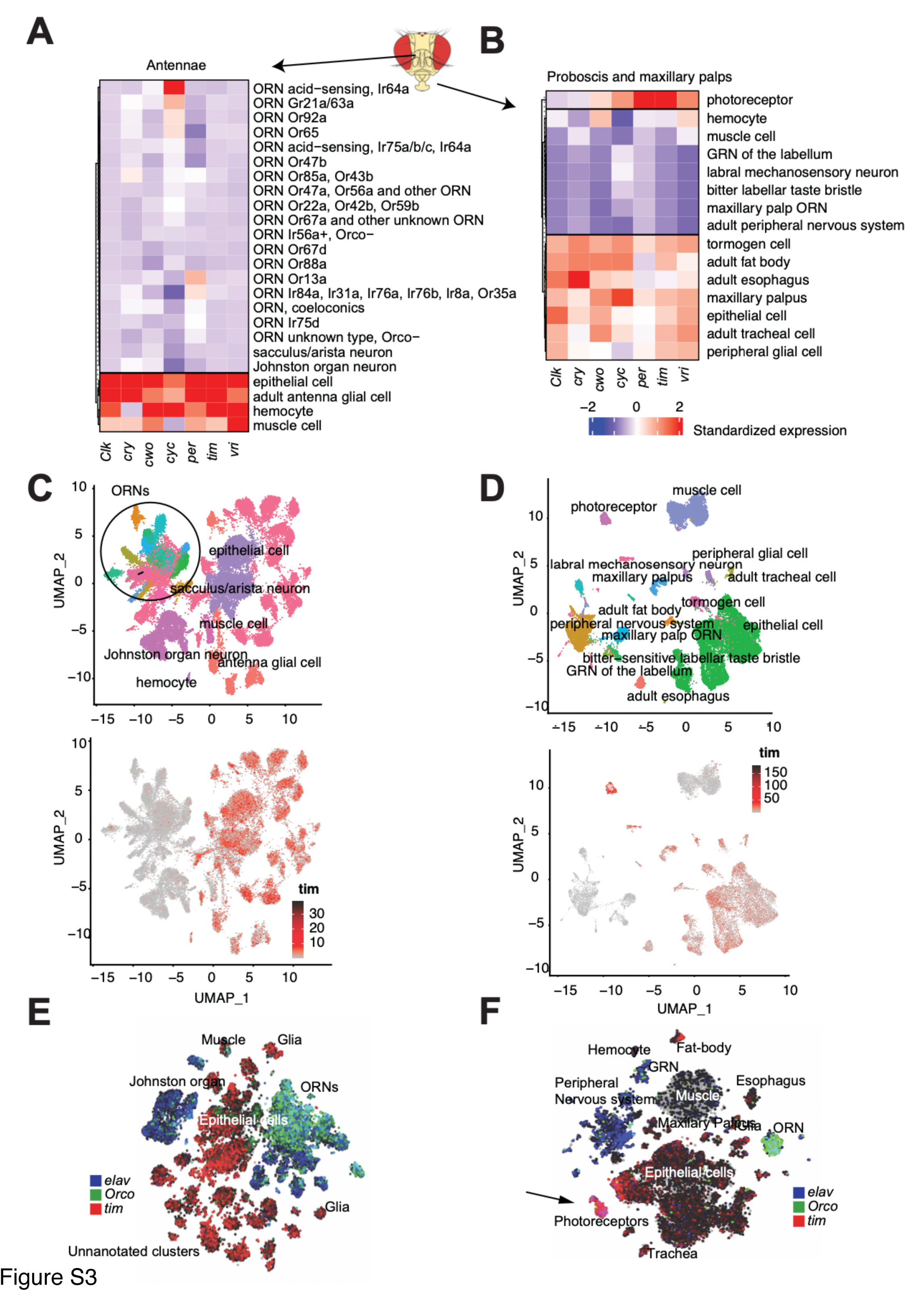
(related to figure 2): Enrichment analysis of core-clock genes in the maxillary antennae (A, C, E) and proboscis and maxillary palp (B, D, F) show low enrichment in GRM and ORM. **A-B:** Heatmaps of standardized values of core-clock genes. **C-D:** UMAP plot showing cluster cell type assignment (top) and normalized *tim* expression (bottom). **E-F:** Co-expression of *elav, Orco,* and *tim* in UMAP plot. In blue *elav* expression; green, *Orco;* and red, *tim.* Overlap of *tim* and *elav* expression is shown in pink (photoreceptors). Light blue marks overlap between *Orco* and *elav* (ORN).

**Figure S4.**
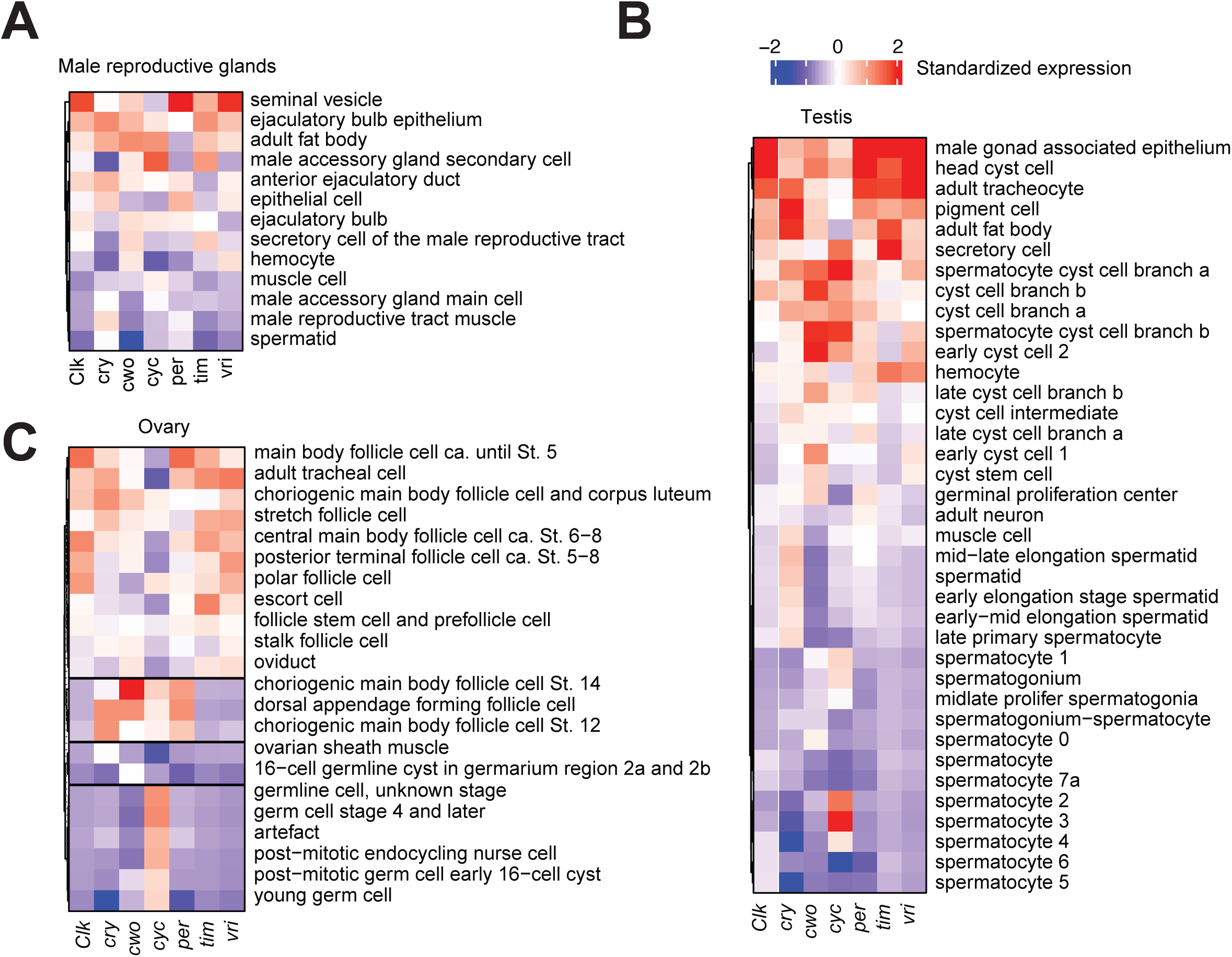
(related to figure 2): Enrichment analysis of core clock genes in testis (A), male reproductive gland (B), and ovary (C) recapitulates previous reports. Heatmaps of standardized values of core-clock for each organ.

**Figure S5.**
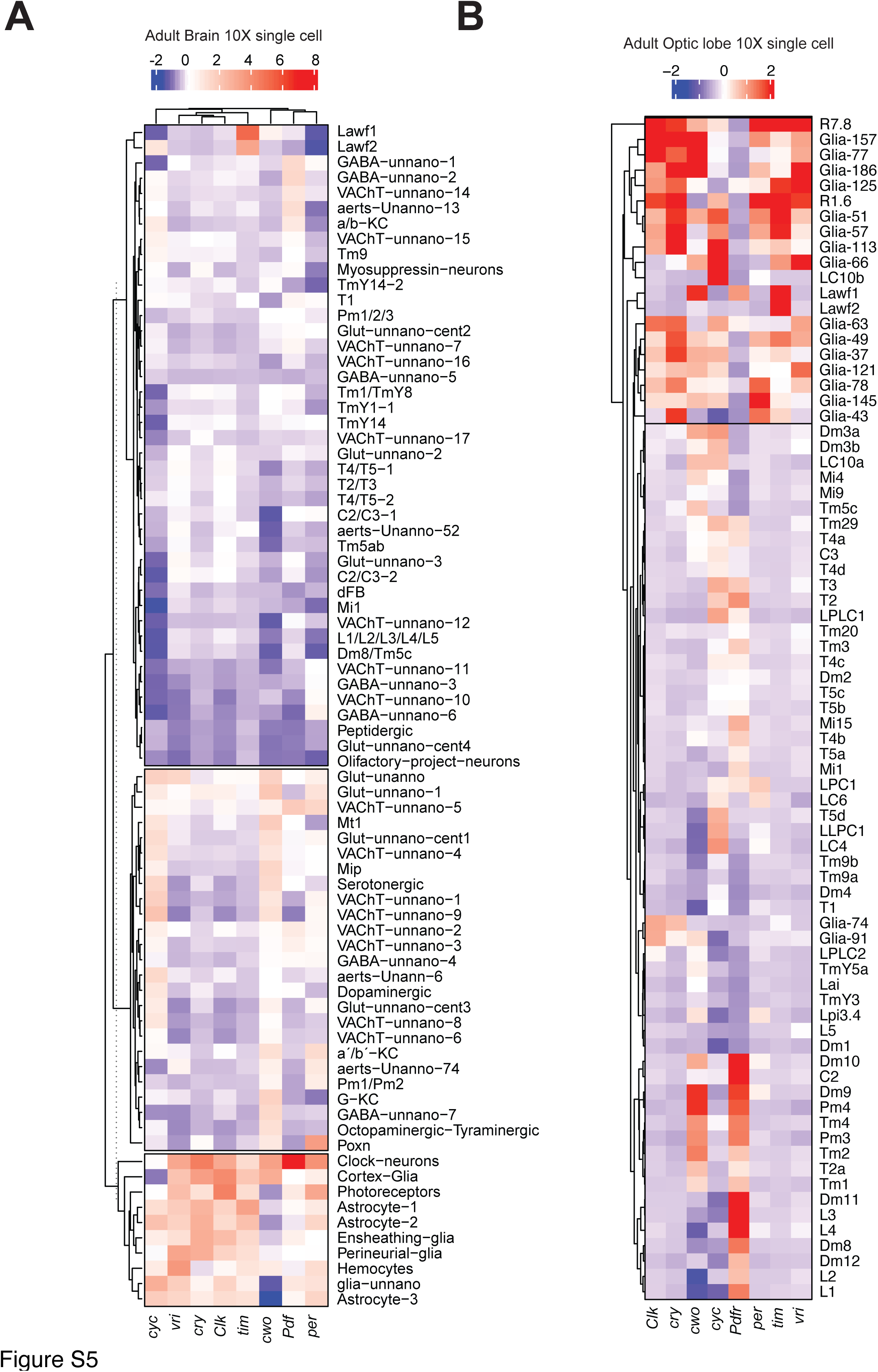
(related to figure 3): Core clock enrichment analysis of 10X single-cell data from the brain (A) and adult optic lobe (B) reveal new cell types expressing high levels of core-clock components. Heatmaps of standardized values of core-clock genes. Blue and red represent low and high levels, respectively.

**Figure S6.**
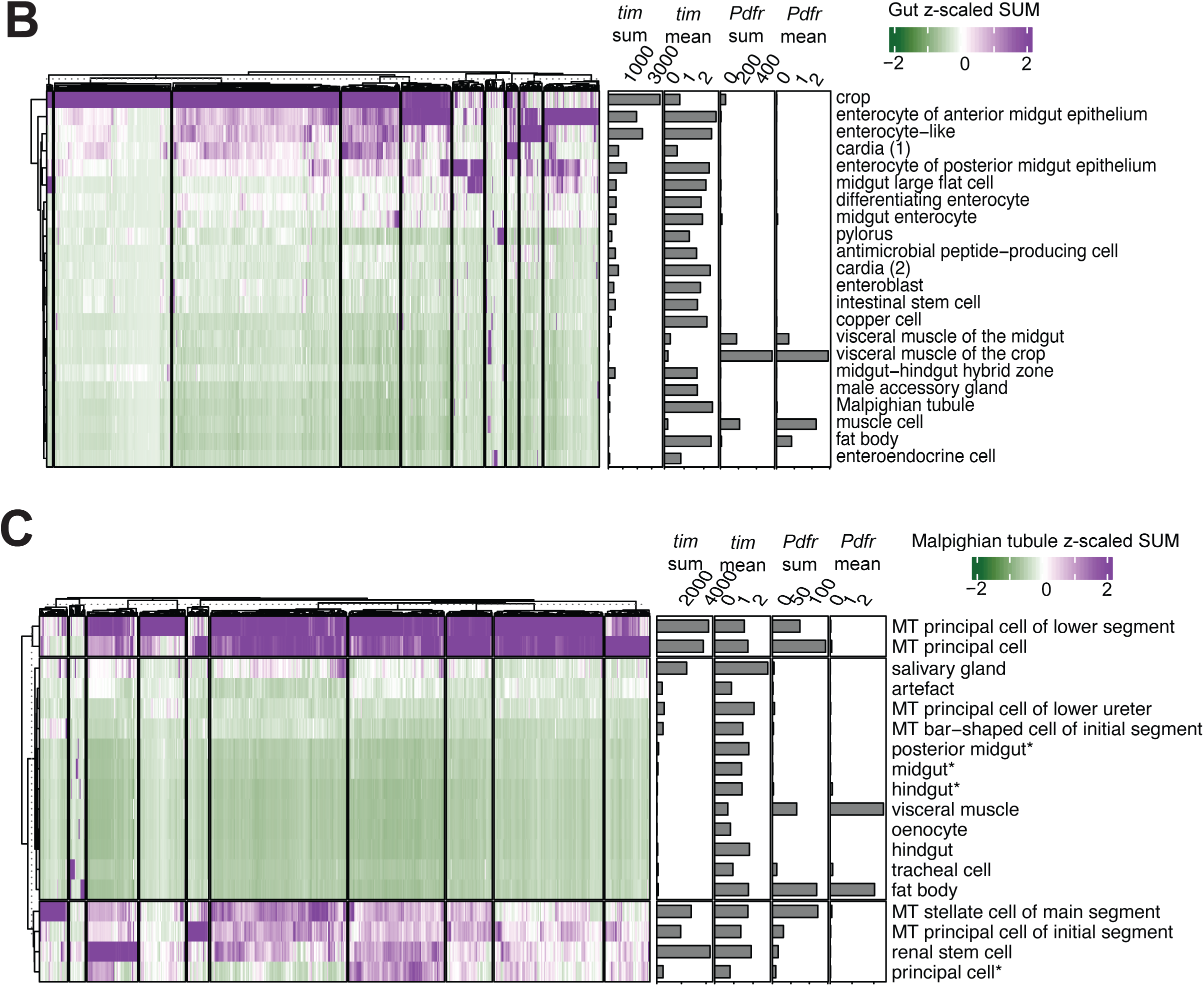
(related to figure 6): Sum contribution of genes with cyclic expression in peripheral tissues reveals enrichment in cells with high levels of *Pdfr* but no *tim*. A. Sum expression of genes cycling in the gut in single-cell clusters from the gut. **B.** Sum expression of Malpighian tubule cyclers single cell clusters. Bar plots represent the mean and sum expression of *tim* and *Pdfr* in each assessed cluster (MT=Malpighian tubule cell)

## REFERENCES

1. Allada R, Chung BY. Circadian organization of behavior and physiology in Drosophila. Annu Rev Physiol. 2010;72:605–24.

2. Pilorz V, Helfrich-Förster C, Oster H. The role of the circadian clock system in physiology. Pflüg Arch – Eur J Physiol. 2018;470(2):227–39.

3. Helfrich-Forster C. The neuroarchitecture of the circadian clock in the brain of Drosophila melanogaster. Microsc Res Tech. 2003;62(2):94–102.

4. Shafer OT, Helfrich-Forster C, Renn SC, Taghert PH. Reevaluation of Drosophila melanogaster’s neuronal circadian pacemakers reveals new neuronal classes. J Comp Neurol. 2006;498(2):180–93.

5. Hardin PE, Panda S. Circadian timekeeping and output mechanisms in animals. Curr Opin Neurobiol. 2013;23(5):724–31.

6. Harms E, Kivimae S, Young MW, Saez L. Posttranscriptional and posttranslational regulation of clock genes. J Biol Rhythms. 2004/11/10 ed. 2004;19(5):361–73.

7. Kojima S, Shingle DL, Green CB. Post-transcriptional control of circadian rhythms. J Cell Sci. 2011/01/19 ed. 2011;124(Pt 3):311–20.

8. Lim C, Allada R. Emerging roles for post-transcriptional regulation in circadian clocks. Nat Neurosci. 2013;16(11):1544–50.

9. Ozkaya O, Rosato E. The circadian clock of the fly: a neurogenetics journey through time. Adv Genet. 2012;77:79–123.

10. Ahmad M, Li W, Top D. Integration of Circadian Clock Information in the Drosophila Circadian Neuronal Network. J Biol Rhythms. 2021 Jun;36(3):203–20.

11. Helfrich-Förster C. Light input pathways to the circadian clock of insects with an emphasis on the fruit fly Drosophila melanogaster. J Comp Physiol A Neuroethol Sens Neural Behav Physiol. 2020;206(2):259–72.

12. Mezan S, Feuz JD, Deplancke B, Kadener S. PDF Signaling Is an Integral Part of the Drosophila Circadian Molecular Oscillator. Cell Rep. 2016/10/13 ed. 2016;17(3):708–19.

13. Peng Y, Stoleru D, Levine JD, Hall JC, Rosbash M. Drosophila free-running rhythms require intercellular communication. PLoS Biol. 2003;1(1):E13–E13.

14. Schlichting M, Menegazzi P, Lelito KR, Yao Z, Buhl E, Dalla Benetta E, et al. A Neural Network Underlying Circadian Entrainment and Photoperiodic Adjustment of Sleep and Activity in Drosophila. J Neurosci. 2016;36(35):9084–96.

15. Stoleru D, Peng Y, Agosto J, Rosbash M. Coupled oscillators control morning and evening locomotor behaviour of Drosophila. Nature. 2004;431(7010):862–8.

16. Weiss R, Bartok O, Mezan S, Malka Y, Kadener S. Synergistic interactions between the molecular and neuronal circadian networks drive robust behavioral circadian rhythms in Drosophila melanogaster. PLoS Genet. 2014/04/05 ed. 2014;10(4):e1004252–e1004252.

17. Klose M, Duvall L, Li W, Liang X, Ren C, Steinbach JH, et al. Functional PDF Signaling in the Drosophila Circadian Neural Circuit Is Gated by Ral A-Dependent Modulation. Neuron. 2016 May 18;90(4):781–94.

18. Im SH, Li W, Taghert PH. PDFR and CRY signaling converge in a subset of clock neurons to modulate the amplitude and phase of circadian behavior in Drosophila. PLoS ONE. 2011;6(4):e18974–e18974.

19. Taghert PH, Shafer OT. Mechanisms of clock output in the Drosophila circadian pacemaker system. J Biol Rhythms. 2006;21(6):445–57.

20. Hyun S, Lee Y, Hong ST, Bang S, Paik D, Kang J, et al. Drosophila GPCR Han is a receptor for the circadian clock neuropeptide PDF. Neuron. 2005;48(2):267–78.

21. Lear BC, Merrill CE, Lin JM, Schroeder A, Zhang L, Allada R. A G protein-coupled receptor, groom-of-PDF, is required for PDF neuron action in circadian behavior. Neuron. 2005;48(2):221–7.

22. Lin Y, Stormo GD, Taghert PH. The neuropeptide pigment-dispersing factor coordinates pacemaker interactions in the Drosophila circadian system. J Neurosci. 2004;24(36):7951– 7.

23. Mertens I, Vandingenen A, Johnson EC, Shafer OT, Li W, Trigg JS, et al. PDF receptor signaling in Drosophila contributes to both circadian and geotactic behaviors. Neuron. 2005;48(2):213–9.

24. Shafer OT, Kim DJ, Dunbar-Yaffe R, Nikolaev VO, Lohse MJ, Taghert PH. Widespread receptivity to neuropeptide PDF throughout the neuronal circadian clock network of Drosophila revealed by real-time cyclic AMP imaging. Neuron. 2008;58(2):223–37.

25. Yoshii T, Wulbeck C, Sehadova H, Veleri S, Bichler D, Stanewsky R, et al. The Neuropeptide Pigment-Dispersing Factor Adjusts Period and Phase of Drosophila’s Clock. J Neurosci. 2009;29(8):2597–610.

26. Zhang L, Chung BY, Lear BC, Kilman VL, Liu Y, Mahesh G, et al. A Subset of Circadian Neurons Coordinates Light and PDF Signaling to Produce Robust Daily Behavior in Drosophila. Curr Biol CB. 2010 Apr 13;20(7):591–9.

27. Hidalgo S, Anguiano M, Tabuloc CA, Chiu JC. Seasonal cues act through the circadian clock and pigment-dispersing factor to control EYES ABSENT and downstream physiological changes. Curr Biol [Internet]. 2023 Jan 27 [cited 2023 Feb 9];0(0). Available from: https://www.cell.com/current-biology/abstract/S0960-9822(23)00006-4

28. He C, Cong X, Zhang R, Wu D, An C, Zhao Z. Regulation of circadian locomotor rhythm by neuropeptide Y-like system in Drosophila melanogaster. Insect Mol Biol. 2013;22(4):376– 88.

29. Johard HAD, Yoishii T, Dircksen H, Cusumano P, Rouyer F, Helfrich-Förster C, et al. Peptidergic clock neurons in Drosophila: ion transport peptide and short neuropeptide F in subsets of dorsal and ventral lateral neurons. J Comp Neurol. 2009 Sep 1;516(1):59–73.

30. Cavanaugh DJ, Geratowski JD, Wooltorton JRA, Spaethling JM, Hector CE, Zheng X, et al. Identification of a circadian output circuit for rest:activity rhythms in Drosophila. Cell. 2014 Apr 24;157(3):689–701.

31. Cavey M, Collins B, Bertet C, Blau J. Circadian rhythms in neuronal activity propagate through output circuits. Nat Neurosci. 2016 Apr;19(4):587–95.

32. Beaver LM, Gvakharia BO, Vollintine TS, Hege DM, Stanewsky R, Giebultowicz JM. Loss of circadian clock function decreases reproductive fitness in males of Drosophila melanogaster. Proc Natl Acad Sci U A. 2002;99(4):2134–9.

33. Chatterjee A, Tanoue S, Houl JH, Hardin PE. Regulation of gustatory physiology and appetitive behavior by the Drosophila circadian clock. Curr Biol. 2010;20(4):300–9.

34. Giebultowicz JM, Stanewsky R, Hall JC, Hege DM. Transplanted Drosophila excretory tubules maintain circadian clock cycling out of phase with the host. Curr Biol. 2000;10(2):107–10.

35. Giebultowicz JM, Ivanchenko M, Vollintine T. – Organization of the insect circadian system: Spatial and developmental expression of clock genes in peripheral tissues of Drosophila melanogaster. In: Denlinger DL, Giebultowicz JM, Saunders DS, editors. Insect Timing: Circadian Rhythmicity to Seasonality [Internet]. Amsterdam: Elsevier Science B.V.; 2001 [cited 2022 Jun 1]. p. 31–42. Available from: https://www.sciencedirect.com/science/article/pii/B9780444506085500350

36. Hege DM, Stanewsky R, Hall JC, Giebultowicz JM. Rhythmic expression of a PER-reporter in the Malpighian tubules of decapitated Drosophila: evidence for a brain-independent circadian clock. J Biol Rhythms. 1997;12(4):300–8.

37. Ito C, Goto SG, Shiga S, Tomioka K, Numata H. Peripheral circadian clock for the cuticle deposition rhythm in Drosophila melanogaster. Proc Natl Acad Sci U A. 2008;105(24):8446–51.

38. Krishnan B, Dryer SE, Hardin PE. Circadian rhythms in olfactory responses of Drosophila melanogaster. Nature. 1999;400(6742):375–8.

39. Levine JD, Funes P, Dowse HB, Hall JC. Advanced analysis of a cryptochrome mutation’s effects on the robustness and phase of molecular cycles in isolated peripheral tissues of Drosophila. BMC Neurosci. 2002;3:5–5.

40. Plautz JD, Kaneko M, Hall JC, Kay SA. Independent photoreceptive circadian clocks throughout Drosophila. Science. 1997;278(5343):1632–5.

41. Myers EM, Yu J, Sehgal A. Circadian control of eclosion: interaction between a central and peripheral clock in Drosophila melanogaster. Curr Biol. 2003;13(6):526–33.

42. Krupp JJ, Kent C, Billeter JC, Azanchi R, So AKC, Schonfeld JA, et al. Social Experience Modifies Pheromone Expression and Mating Behavior in Male Drosophila melanogaster. Curr Biol. 2008 Sep 23;18(18):1373–83.

43. Sakai T, Ishida N. Circadian rhythms of female mating activity governed by clock genes in Drosophila. Proc Natl Acad Sci U A. 2001;98(16):9221–5.

44. Tanoue S, Krishnan P, Krishnan B, Dryer SE, Hardin PE. Circadian clocks in antennal neurons are necessary and sufficient for olfaction rhythms in Drosophila. Curr Biol. 2004;14(8):638–49.

45. Krishnan B, Levine JD, Lynch MK, Dowse HB, Funes P, Hall JC, et al. A new role for cryptochrome in a Drosophila circadian oscillator. Nature. 2001/05/18 ed. 2001;411(6835):313–7.

46. Krupp JJ, Billeter JC, Wong A, Choi C, Nitabach MN, Levine JD. Pigment-Dispersing Factor Modulates Pheromone Production in Clock Cells that Influence Mating in Drosophila. Neuron. 2013 Jul 10;79(1):54–68.

47. Versteven M, Ernst KM, Stanewsky R. A Robust and Self-Sustained Peripheral Circadian Oscillator Reveals Differences in Temperature Compensation Properties with Central Brain Clocks. iScience. 2020 Aug 21;23(8):101388.

48. Litovchenko M, Meireles-Filho ACA, Frochaux MV, Bevers RPJ, Prunotto A, Anduaga AM, et al. Extensive tissue-specific expression variation and novel regulators underlying circadian behavior. Sci Adv. 2021 Jan 29;7(5):eabc3781.

49. Xu K, DiAngelo JR, Hughes ME, Hogenesch JB, Sehgal A. The Circadian Clock Interacts with Metabolic Physiology to Influence Reproductive Fitness. Cell Metab. 2011 Jun 8;13(6):639–54.

50. Zhang Y, Li Y, Barber AF, Noya SB, Williams JA, Li F, et al. The microbiome stabilizes circadian rhythms in the gut. Proc Natl Acad Sci. 2023 Jan 31;120(5):e2217532120.

51. Erion R, King AN, Wu G, Hogenesch JB, Sehgal A. Neural clocks and Neuropeptide F/Y regulate circadian gene expression in a peripheral metabolic tissue. Vijay Raghavan K, editor. eLife. 2016 Apr 14;5:e13552.

52. Chen DM, Christianson JS, Sapp RJ, Stark WS. Visual receptor cycle in normal and period mutant Drosophila: Microspectrophotometry, electrophysiology, and ultrastructural morphometry. Vis Neurosci. 1992 Aug;9(2):125–35.

53. Ogueta M, Hardie RC, Stanewsky R. Non-canonical Phototransduction Mediates Synchronization of the Drosophila melanogaster Circadian Clock and Retinal Light Responses. Curr Biol. 2018/05/22 ed. 2018;28(11):1725–1735 e3.

54. Damulewicz M, Rosato E, Pyza E. Circadian regulation of the na(+)/k(+)-ATPase alpha subunit in the visual system is mediated by the pacemaker and by retina photoreceptors in Drosophila melanogaster. PLoS ONE. 2013;8(9):e73690–e73690.

55. Górska-Andrzejak J, Keller A, Raabe T, Kilianek L, Pyza E. Structural daily rhythms in GFP-labelled neurons in the visual system of Drosophila melanogaster. Photochem Photobiol Sci Off J Eur Photochem Assoc Eur Soc Photobiol. 2005 Sep;4(9):721–6.

56. Pyza E, Górska-Andrzejak J. Involvement of glial cells in rhythmic size changes in neurons of the housefly’s visual system. J Neurobiol. 2004;59(2):205–15.

57. Pyza E, Meinertzhagen IA. Daily rhythmic changes of cell size and shape in the first optic neuropil in Drosophila melanogaster. J Neurobiol. 1999 Jul;40(1):77–88.

58. Weber P, Kula-Eversole E, Pyza E. Circadian control of dendrite morphology in the visual system of Drosophila melanogaster. PLoS ONE. 2009;4(1):e4290–e4290.

59. Damulewicz M, Loboda A, Bukowska-Strakova K, Jozkowicz A, Dulak J, Pyza E. Clock and clock-controlled genes are differently expressed in the retina, lamina and in selected cells of the visual system of Drosophila melanogaster. Front Cell Neurosci [Internet]. 2015 [cited 2022 Jul 8];9. Available from: https://www.frontiersin.org/articles/10.3389/fncel.2015.00353

60. Davie K, Janssens J, Koldere D, De Waegeneer M, Pech U, Kreft L, et al. A Single-Cell Transcriptome Atlas of the Aging Drosophila Brain. Cell. 2018/06/19 ed. 2018;174(4):982–998 e20.

61. Konstantinides N, Kapuralin K, Fadil C, Barboza L, Satija R, Desplan C. Phenotypic Convergence: Distinct Transcription Factors Regulate Common Terminal Features. Cell. 2018/06/19 ed. 2018;174(3):622–635 e13.

62. Kurmangaliyev YZ, Yoo J, Valdes-Aleman J, Sanfilippo P, Zipursky SL. Transcriptional Programs of Circuit Assembly in the Drosophila Visual System. Neuron. 2020 Dec 23;108(6):1045–1057.e6.

63. Li H, Janssens J, De Waegeneer M, Kolluru SS, Davie K, Gardeux V, et al. Fly Cell Atlas: A single-nucleus transcriptomic atlas of the adult fruit fly. Science. 2022 Mar 4;375(6584):eabk2432.

64. Gunawardhana KL, Rivas GBS, Caster C, Hardin PE. Crosstalk between vrille transcripts, proteins, and regulatory elements controlling circadian rhythms and development in Drosophila. iScience [Internet]. 2021 Jan 22 [cited 2022 Jul 10];24(1). Available from: https://www.cell.com/iscience/abstract/S2589-0042(20)31090-7

65. Iyer EPR, Iyer SC, Sullivan L, Wang D, Meduri R, Graybeal LL, et al. Functional Genomic Analyses of Two Morphologically Distinct Classes of Drosophila Sensory Neurons: Post-Mitotic Roles of Transcription Factors in Dendritic Patterning. PLOS ONE. 2013 Aug 15;8(8):e72434.

66. Kadener S, Stoleru D, McDonald M, Nawathean P, Rosbash M. Clockwork Orange is a transcriptional repressor and a new Drosophila circadian pacemaker component. Genes Dev. 2007/06/21 ed. 2007;21(13):1675–86.

67. Szuplewski S, Fraisse-Véron I, George H, Terracol R. vrille is required to ensure tracheal integrity in Drosophila embryo. Dev Growth Differ. 2010;52(5):409–18.

68. Liu T, Mahesh G, Yu W, Hardin PE. CLOCK stabilizes CYCLE to initiate clock function in Drosophila. Proc Natl Acad Sci. 2017 Oct 10;114(41):10972–7.

69. Davis FP, Nern A, Picard S, Reiser MB, Rubin GM, Eddy SR, et al. A genetic, genomic, and computational resource for exploring neural circuit function. Bellen HJ, Vijay Raghavan K, editors. eLife. 2020 Jan 15;9:e50901.

70. Zappia MP, Rogers A, Islam ABMMK, Frolov MV. Rbf Activates the Myogenic Transcriptional Program to Promote Skeletal Muscle Differentiation. Cell Rep. 2019 Jan 15;26(3):702–719.e6.

71. Chatterjee A, Hardin PE. Time to taste: circadian clock function in the Drosophila gustatory system. Fly Austin. 2010;4(4):283–7.

72. Beaver LM, Rush BL, Gvakharia BO, Giebultowicz JM. Noncircadian regulation and function of clock genes period and timeless in oogenesis of Drosophila melanogaster. J Biol Rhythms. 2003 Dec;18(6):463–72.

73. Krzeptowski W, Walkowicz L, Płonczyńska A, Górska-Andrzejak J. Different Levels of Expression of the Clock Protein PER and the Glial Marker REPO in Ensheathing and Astrocyte-Like Glia of the Distal Medulla of Drosophila Optic Lobe. Front Physiol [Internet]. 2018 [cited 2022 Jun 1];9. Available from: https://www.frontiersin.org/article/10.3389/fphys.2018.00361

74. You S, Yu AM, Roberts MA, Joseph IJ, Jackson FR. Circadian regulation of the Drosophila astrocyte transcriptome. PLoS Genet. 2021 Sep;17(9):e1009790.

75. Jauregui-Lozano J, Hall H, Stanhope SC, Bakhle K, Marlin MM, Weake VM. The Clock:Cycle complex is a major transcriptional regulator of Drosophila photoreceptors that protects the eye from retinal degeneration and oxidative stress. PLOS Genet. 2022 Jan 31;18(1):e1010021.

76. Shirasu-Hiza MM, Dionne MS, Pham LN, Ayres JS, Schneider DS. Interactions between circadian rhythm and immunity in Drosophila melanogaster. Curr Biol. 2007;17(10):R353–5.

77. Chen Z, Del Valle Rodriguez A, Li X, Erclik T, Fernandes VM, Desplan C. A Unique Class of Neural Progenitors in the Drosophila Optic Lobe Generates Both Migrating Neurons and Glia. Cell Rep. 2016 Apr 26;15(4):774–86.

78. Tuthill JC, Nern A, Rubin GM, Reiser MB. Wide-field feedback neurons dynamically tune early visual processing. Neuron. 2014 May 21;82(4):887–95.

79. Yuan D, Ji X, Hao S, Gestrich JY, Duan W, Wang X, et al. Lamina feedback neurons regulate the bandpass property of the flicker-induced orientation response in Drosophila. J Neurochem. 2021;156(1):59–75.

80. Tuthill JC, Nern A, Holtz SL, Rubin GM, Reiser MB. Contributions of the 12 Neuron Classes in the Fly Lamina to Motion Vision. Neuron. 2013 Jul 10;79(1):128–40.

81. Abruzzi KC, Zadina A, Luo W, Wiyanto E, Rahman R, Guo F, et al. RNA-seq analysis of Drosophila clock and non-clock neurons reveals neuron-specific cycling and novel candidate neuropeptides. PLoS Genet. 2017/02/10 ed. 2017;13(2):e1006613–e1006613.

82. Duffield GE, Best JD, Meurers BH, Bittner A, Loros JJ, Dunlap JC. Circadian programs of transcriptional activation, signaling, and protein turnover revealed by microarray analysis of mammalian cells. Curr Biol CB. 2002 Apr 2;12(7):551–7.

83. Hughes ME, Grant GR, Paquin C, Qian J, Nitabach MN. Deep sequencing the circadian and diurnal transcriptome of Drosophila brain. Genome Res. 2012/04/05 ed. 2012;22(7):1266–81.

84. Kuintzle RC, Chow ES, Westby TN, Gvakharia BO, Giebultowicz JM, Hendrix DA. Circadian deep sequencing reveals stress-response genes that adopt robust rhythmic expression during aging. Nat Commun. 2017 Feb 21;8(1):14529.

85. Martin Anduaga A, Evantal N, Patop IL, Bartok O, Weiss R, Kadener S. Thermosensitive alternative splicing senses and mediates temperature adaptation in Drosophila. Ramaswami M, Calabrese RL, editors. eLife. 2019 Nov 8;8:e44642.

86. Rodriguez J, Tang CH, Khodor YL, Vodala S, Menet JS, Rosbash M. Nascent-Seq analysis of Drosophila cycling gene expression. Proc Natl Acad Sci U A. 2013/01/09 ed. 2013;110(4):E275–84.

87. Casey T, Suarez-Trujillo A, Cummings S, Huff K, Crodian J, Bhide K, et al. Core circadian clock transcription factor BMAL1 regulates mammary epithelial cell growth, differentiation, and milk component synthesis. PLOS ONE. 2021 Aug 20;16(8):e0248199.

88. Gibbs JE, Beesley S, Plumb J, Singh D, Farrow S, Ray DW, et al. Circadian timing in the lung; a specific role for bronchiolar epithelial cells. Endocrinology. 2009 Jan;150(1):268– 76.

89. Keller M, Mazuch J, Abraham U, Eom GD, Herzog ED, Volk HD, et al. A circadian clock in macrophages controls inflammatory immune responses. Proc Natl Acad Sci. 2009 Dec 15;106(50):21407–12.

90. Sládek M, Rybová M, Jindráková Z, Zemanová Z, Polidarová L, Mrnka L, et al. Insight Into the Circadian Clock Within Rat Colonic Epithelial Cells. Gastroenterology. 2007 Oct 1;133(4):1240–9.

91. Williams J, Yang N, Wood A, Zindy E, Meng QJ, Streuli CH. Epithelial and stromal circadian clocks are inversely regulated by their mechano-matrix environment. J Cell Sci. 2018 Mar 1;131(5):jcs208223.

92. Park WS, Miyano-Kurosaki N, Abe T, Takai K, Yamamoto N, Takaku H. Inhibition of HIV-1 replication by a new type of circular dumbbell RNA/DNA chimeric oligonucleotides. Biochem Biophys Res Commun. 2000;270(3):953–60.

93. Im SH, Taghert PH. PDF receptor expression reveals direct interactions between circadian oscillators in Drosophila. J Comp Neurol. 2010;518(11):1925–45.

94. Talsma AD, Christov CP, Terriente-Felix A, Linneweber GA, Perea D, Wayland M, et al. Remote control of renal physiology by the intestinal neuropeptide pigment-dispersing factor in Drosophila. Proc Natl Acad Sci. 2012 Jul 24;109(30):12177–82.

95. Hermann C, Yoshii T, Dusik V, Helfrich-Forster C. Neuropeptide F immunoreactive clock neurons modify evening locomotor activity and free-running period in Drosophila melanogaster. J Comp Neurol. 2011/08/10 ed. 2012;520(5):970–87.

96. Selcho M, Millán C, Palacios-Muñoz A, Ruf F, Ubillo L, Chen J, et al. Central and peripheral clocks are coupled by a neuropeptide pathway in Drosophila. Nat Commun. 2017 May 30;8(1):15563.

97. Hashimshony T, Wagner F, Sher N, Yanai I. CEL-Seq: single-cell RNA-Seq by multiplexed linear amplification. Cell Rep. 2012;2(3):666–73.

98. Hashimshony T, Senderovich N, Avital G, Klochendler A, de Leeuw Y, Anavy L, et al. CEL-Seq2: sensitive highly-multiplexed single-cell RNA-Seq. Genome Biol. 2016/04/29 ed. 2016;17:77–77.

99. Dobin A, Davis CA, Schlesinger F, Drenkow J, Zaleski C, Jha S, et al. STAR: ultrafast universal RNA-seq aligner. Bioinformatics. 2013;29(1):15–21.

100. Derr A, Yang C, Zilionis R, Sergushichev A, Blodgett DM, Redick S, et al. End Sequence Analysis Toolkit (ESAT) expands the extractable information from single-cell RNA-seq data. Genome Res. 2016;26(10):1397–410.

101. Wu G, Anafi RC, Hughes ME, Kornacker K, Hogenesch JB. MetaCycle: an integrated R package to evaluate periodicity in large scale data. Bioinformatics. 2016/10/30 ed. 2016;32(21):3351–3.

102. Hughes ME, Abruzzi KC, Allada R, Anafi R, Arpat AB, Asher G, et al. Guidelines for Genome-Scale Analysis of Biological Rhythms. J Biol Rhythms. 2017/11/04 ed. 2017;32(5):380–93.

